# UCA1 lncRNA represses γ-globin expression by sequestering miR-148b, a key post-transcriptional regulator of BCL11A

**DOI:** 10.1101/2025.08.29.673129

**Authors:** Motiur Rahaman, Shatarupa Bhattacharya, Mandrita Mukherjee, Chiranjib Bhowmick, Praphulla Chandra Shukla, Tuphan Kanti Dolai, Nishant Chakravorty

**Affiliations:** School of Medical Science and Technology, IIT Kharagpur, Kharagpur, Paschim Medinipur, West Bengal – 721302, India; Department of Hematology, Nil Ratan Sircar Medical College and Hospital, Kolkata, West Bengal-700014, India

**Keywords:** LncRNA, miRNA, ceRNA network, Fetal hemoglobin, β-hemoglobinopathies

## Abstract

Fetal hemoglobin (HbF; α₂γ₂) reactivation is a promising strategy to ameliorate β-hemoglobinopathies. However, limited understanding of γ-globin (HBG1/2) regulation constrains development of therapeutic interventions. BCL11A, a key transcriptional repressor of γ-globin, is central to HbF silencing during adult erythropoiesis. Here, we identify a new post-transcriptional regulatory mechanism involving the lncRNA-UCA1 and miR-148b that modulates BCL11A expression. Using UCA1 knockdown and overexpression strategies, combined with in vivo crosslinking and transcriptomic analyses, we demonstrate that UCA1 functions as a competing endogenous RNA (ceRNA), sequestering miR-148b and thereby attenuating its repressive effect on BCL11A. In the present study, we have elucidated the physiological significance of this interaction in adult erythroid cells, including CD34⁺ HSPCs and HUDEP-2 cells, in which UCA1 depletion led to robust γ-globin induction, a phenotype recapitulated by miR-148b overexpression. These findings uncover a previously unrecognized lncRNA-miRNA-mRNA regulatory axis and highlight the UCA1/miR-148b axis as a potential therapeutic target for HbF reactivation in β-hemoglobinopathies.

## INTRODUCTION

Fetal hemoglobin (HbF, α_2_γ_2_) is the predominant form of hemoglobin at birth. The switch from γ-globin to β-globin gene expression occurs during ontogeny, leading to the gradual replacement of fetal hemoglobin (HbF) by adult hemoglobin (HbA)[1]. The complex regulatory mechanisms underlying globin gene switching are crucial to our understanding of higher eukaryotic transcriptional control. In healthy adults, HbF typically constitutes less than 1% of total hemoglobin and has limited physiological relevance[2]. However, congenital, acquired, and drug-inductionstrategies for HbF induction are reported to ameliorate the clinical manifestations of β-hemoglobinopathies, such as Sickle cell anemia (SCA) and β-thalassemia, by reducing the α-globin chain precipitation and providing an alternate form of functional hemoglobin. Therefore, unraveling the mechanisms of γ-globin gene regulation holds direct translational relevance for developing newtherapies in β-hemoglobinopathies[3]. Both genetic and epigenetic factors are known to influence γ-globin expression in adult erythroid cells. Naturally occurring conditions such as hereditary persistence of fetal hemoglobin (HPFH), δβ-thalassemia, and pathological disorders like juvenile myelomonocytic leukemia (JMML) offer valuable insights into adult γ-globin regulation[4][5]. Nevertheless, a detailed understanding regarding the mechanisms to elevate γ-globin expression remains incomplete and continues to attract considerable scientific attention.

As efforts to elevate HbF levels intensify, growing evidence highlights the critical role of non-coding RNAs (ncRNAs) in modulating globin gene expression. These regulatory RNAs offer promising strategies for reactivating γ-globin genes in adult erythroid cells[6]. Among them, long non-coding RNAs (lncRNAs) - a major class of non-coding RNAs (ncRNAs) with transcript length longer than 200 nucleotides, have emerged as major focus of translational research in the recent decades. Once dismissed as transcriptional noise, lncRNAs are now recognized for their regulatory functions at transcriptional, post-transcriptional, and translational levels. They modulate processes such as chromatin remodeling, nuclear architecture, cellular differentiation, and gene expression[7]. Over the past decade, numerous lncRNAs have been identified in erythroid cells, with several shown to influence erythropoiesis and red blood cell maturation[8]. Despite widespread abundance and functional relevance, only a few erythroid-specific lncRNAs have been characterized so far.

Another important class of ncRNAs are microRNAs (miRNAs), small ∼22 nucleotide transcripts, which regulate gene expression post-transcriptionally through transcript degradation or translation inhibition[9].Each target mRNA transcript which shares the same miRNA response elements(MREs), can influence the specific cellular functions by participating in the competing endogenous RNA networks (ceRNA) for common miRNA repertoire[10].

Previously, it has been reported that lncRNAs can elicit their function by acting as miRNA decoys or sponges and can affect different biological activities by modulating the availability of miRNAs across their target transcripts[11]. This competitive binding may occur through direct interaction with the miRNA itself or with the 3′ untranslated region (3′UTR) of target mRNAs, thereby mitigating miRNA-mediated silencing[12][13]. Additionally, lncRNAs have been shown to act as precursors of miRNAs and can regulate miRNA biogenesis in different pathophysiological contexts [14]. Recent studies have uncovered emerging roles for lncRNAs in HbF regulation, providing cues for HbF reactivation through non-coding RNA-mediated networks[15][16]. It is expected that identification of new lncRNAs would contribute substantially to our understanding of the regulatory mechanisms of the γ-globin genes and hence bring us closer to developing new therapeutic strategies to treat β-hemoglobinopathies. Genome-wide association studies (GWAS) have identified three key loci-HBB, the HBS1L-MYB intergenic region, and BCL11A, that collectively account for approximately 20–45% of HbF variation across populations [17]. In the HBS1L-MYB locus, a 3-bp deletion polymorphism within an enhancer gives rise to a 1283-bp transcript termed HMI-lncRNA. Knockdown of this lncRNA results in a dramatic (∼200-fold) increase in γ-globin expression. Although the precise binding targets and downstream pathways remain unclear, HMI-lncRNA is hypothesized to interact with the MYB promoter, thereby modulating HbF levels[18]. These findings underscore the crucial role of lncRNAs in orchestrating fetal-to-adult globin switching.

Despite the discovery of a repertoire of lncRNAs, the functional significance of lncRNA-mediated sponges in γ-globin gene regulation remains largely unexplored. The present study systematically investigates the lncRNA-mediated sponge regulatory network inγ-globin genes expression. By integrating sequence-based prediction with expression profiling in adult erythroid cells, we identified UCA1 as a putative miR-148b sponge. Functional validation experiments revealed that UCA1 negatively regulates γ-globin expression via miR-148b sequestration. To our knowledge, this is the first demonstration of UCA1 acting as a miRNA sponge to modulate γ-globin expression. Our findings provide mechanistic insight into a novel lncRNA–miRNA–γ-globin axis and suggest that modulation of lncRNA-mediated sponge activity may represent a promising therapeutic strategy for HbF induction in β-hemoglobinopathies.

## RESULTS

### In-silico prediction of sponge lncRNAs regulating γ-Globin-associated genes

Sponge long non-coding RNAs (splncRNAs) are a distinct class of gene expression regulators that, unlike transcription factors, exert their function by sharing common microRNA (miRNA) binding sites, thereby modulating the availability of miRNAs to target mRNAs. The efficacy of this interaction is influenced by the intracellular stoichiometry of miRNAs and their competing RNA targets.

To identify splncRNAs potentially regulating γ-globin-associated genes, we developed a computational pipeline to extract lncRNA signatures from microarray data by repurposing Affymetrix Exon Array probes. Although these datasets were not originally designed to capture lncRNA expression, valuable information can be retrieved through probe re-annotation. Using RefSeq and Ensembl annotations, we repurposed the probes and identified a total of 2,248 unique lncRNAs. Differential expression analysis revealed 37 lncRNAs (20 up-regulated, 17 down-regulated) significantly associated with fetal hemoglobin (HbF) regulation (Figure 1A). The complete list of 37 differentially expressed lncRNAs is presented in Supplemental information (Table S2). These 37 lncRNAs were subjected to further feature selection and classification, using a supervised machine learning approach. Specifically, we employed a Support Vector Machine (SVM) with a radial basis function (RBF) kernel alongside a stepwise feature selection method.

**Figure 1.**
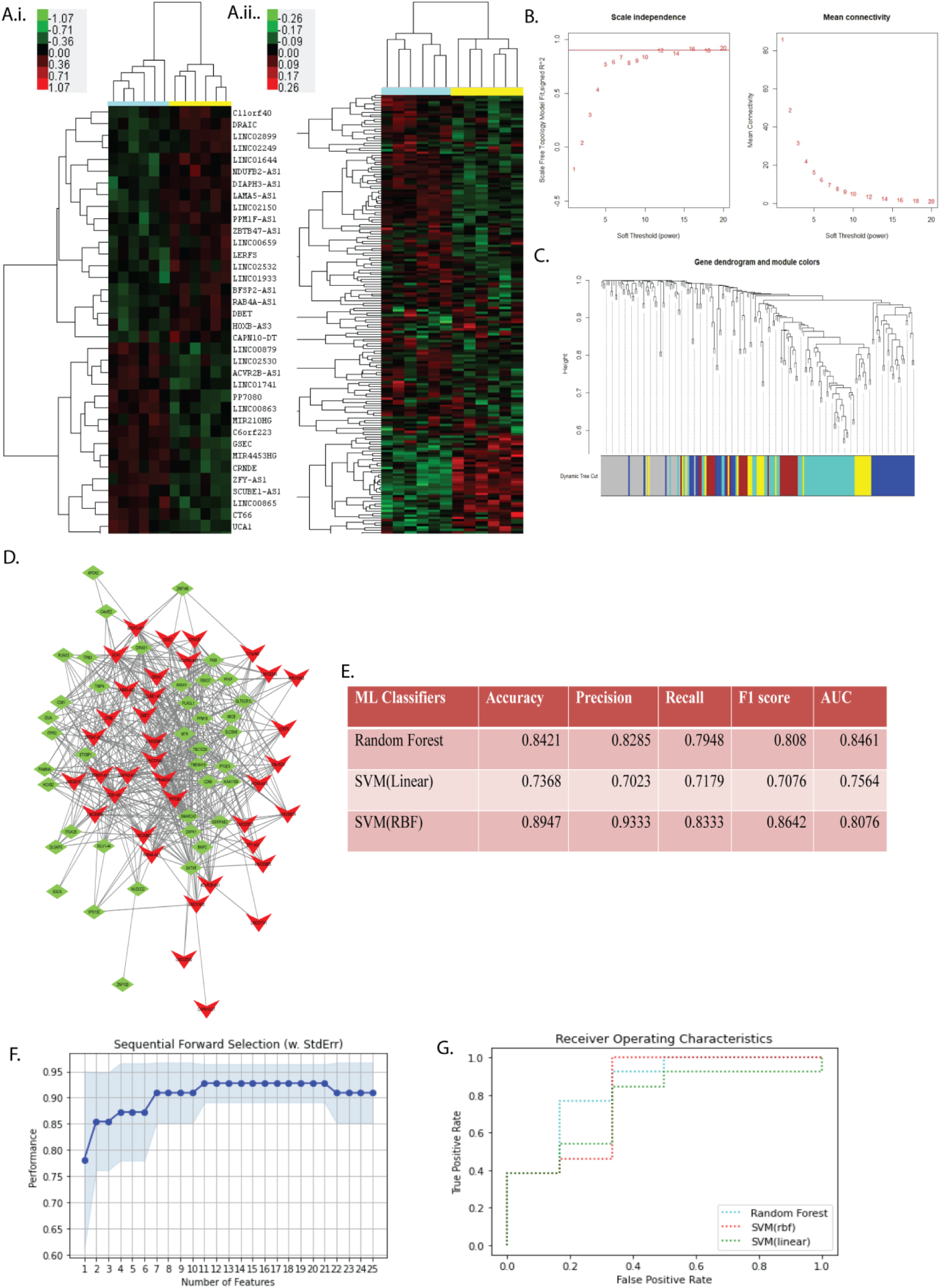
Integrated computational pipeline for identifying lncRNAs associated with γ-globin regulation. (A.i) Heat map showing differentially expressed lncRNAs between high HbF and normal condition. (A.ii) Heat map showing differentially expressed genes between high HbF and normal HbF condition. Red= high expression; Green=low expression. Bar colours indicate sample type: yellow, high HbF; blue, normal HbF. (B.) WCGNA analysis plots. Scale independence and mean connectivity analysis. (C.) Figure showing clustering dendogram of genes obtained by using adjacent dissimilarity-based hierarchical clustering. (D.)Representative colour module, blue network module is shown. The network analysis indicates that highest score is related to UCA1. (E.) Figure is showing performance matrix of different Machine Learning (ML) classifiers. (F.) Showing features selection procedure to identify optimal set of lncRNA from differentially expressed lncRNAs. Stepwise selection method, involving forward selection was used. (G.) Figure is showing ROC curve of lncRNA-based classifiers in the discovery cohort.

This analysis identified 11 lncRNAs (C6orf223, AP000679.1, GSEC, UCA1, CT66, MIR4453HG, AL135925.1, ZEB1-AS1, AC091057.1, PPM1F-AS1, and LINC02249) as the optimal combination for predicting high HbF conditions. To assess predictive performance, we utilized three different classifiers-Random Forest, SVM with linear kernel (SVM-L), and SVM with RBF kernel (SVM-RBF). Among them, the SVM-RBF classifier exhibited the highest performance, achieving an accuracy of 89.47% and a precision of 93.33% (Figure 1E).

### Identification of novel lncRNA modules

To construct lncRNA-based gene regulatory networks associated with γ-globin expression, we performed Weighted Gene Co-expression Network Analysis (WGCNA). This analysis revealed several modules of co-expressed genes, including novel lncRNAs tightly correlated with differentially expressed protein-coding genes involved in fetal hemoglobin (HbF) regulation. Notably, key lncRNAs such as UCA1, GSEC, ZEB1-AS1, and MIR4453HG were consistently enriched across major color-coded modules (blue, brown, turquoise, and yellow), except the grey module. Among them, UCA1 emerged as a central hub with the highest number of predicted interactions, suggesting its prominent role as a potential sponge regulator in the γ-globin regulatory network. Interaction networks were constructed using the CytoHubba plugin in Cytoscape (Figure 1D).The integration of WGCNA and network topology analyses further supports the functional relevance of UCA1 and other splncRNAs in modulating γ-globin expression through competitive endogenous RNA (ceRNA) network.

### Experimental validation of lncRNA-based regulatory networks in an *in-vitro* model of high HbF

To validate the predicted lncRNA-mediated regulatory networks, we developed an in vitro model mimicking the high HbF condition using K562,a human erythroleukemic cell line. The γ-globin expression was induced by treating the erythroid differentiated cells with Hydroxyurea (HU). In order to assess erythroid differentiation, we performed RT-qPCR analysis of marker genes (BAND3, GATA1, and ALAS2) at different time points, which confirmed a steady increase in their expressions (Figure 2D).Erythroid differentiation was further validated by Giemsa-Wright staining and flow cytometry, with 78.7% of HU-treated cells exhibiting a CD71⁺/CD235⁺ phenotype at 72h, compared to 56.6% in untreated cells (Figures 2A-C).Next, we aimed to mimic a high HbF condition by treating the cells with Hydroxyurea (HU) and assess γ-globin expression.

**Figure 2.**
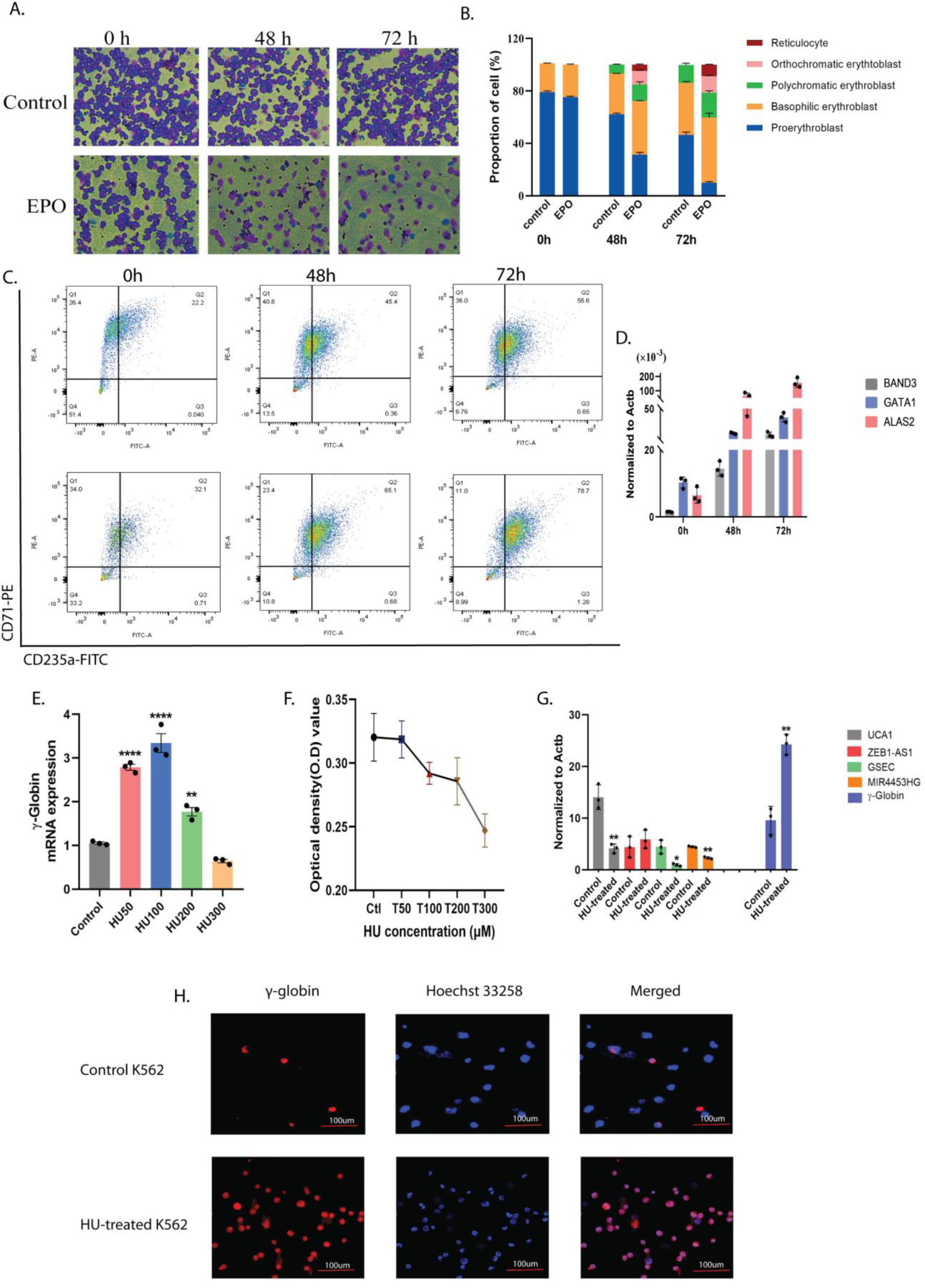
Experimental validation of key lncRNAs predicted from *in-silico* analysis. (A.) Representative staining images showing morphological changes of K562 cells post EPO (3IU/ml) treatment. (B.) Bar plots showing different proportions of erythroid lineage cells post EPO (3IU/ml) treatment. (C.) Figure showing different stages of erythroid cells post EPO treatment. Representative FACS plots showing different proportions of CD71+/CD235a+ erythroid cells at different time points (0-72h) post EPO (3IU/ml) treatment. (D.) RT-qPCR analysis showing expression levels of key markers of erythroid differentiation, BAND3, GATA1, and ALAS2. (E.) RT-qPCR analysis showing expression levels of γ-globin in K562 cells treated with increasing concentrations (50-300µM) of Hydroxyurea (HU) drug. (F.) Figure showing the MTT assay results in K562 cells after HU treatments. The x-axis represents drug concentrations, while the y-axis represents absorbance value obtained from MTT assay. (G.) RT-qPCR analysis showing expression of different predicted lncRNAs in K562 cells post HU-treatment in comparison to non-treated control. (H.) Fluorescence image showing up-regulation of γ-globin post HU-treatment in comparison to non-treated control. The scale bar indicates 100µm. Data are representative of three independent biological replicates and graphical data are represented as Mean ± SD. * p <0.05, ** p <0.01, **** p<0.0001 by unpaired two-tailed t-test.

To optimize HU dosage, cells were treated with increasing concentrations (50–300 µM) of the HU-drug. Flow cytometric analysis of F-cell staining revealed a dose-dependent increase in HbF levels with HU-treatment (Supplemental Figure S3.B).RT-qPCR analysis at 24 h post treatment showed a dose-dependent increase in γ-globin expression, peaking at 100 µM, after which expression declined (Figure 2E). Cell viability also dropped significantly beyond 100 µM HU (Figure 2F), establishing 100 µM as the optimal dose. Immunoblot analysis followed by Immunofluorescence study confirmed a marked increase in γ-globin expression at this dose (Supplemental Figure S3.E and Figure 2H).Once the high HbF condition was established, we proceeded to validate the expression of the predicted lncRNAs identified through our in silico analysis. RT-qPCR analysis of predicted lncRNAs revealed significant down-regulation of UCA1, GSEC, and MIR4453HG in high HbF conditions, consistent with our in-silico predictions (Figure 2G). While ZEB1-AS1 was predicted to be down-regulated, no significant change was observed experimentally. Given UCA1 showed high network connectivity and consistent down-regulation, it was prioritized for further sponge network construction and functional analysis.

### Coordinated expression of UCA1 and miR-148b up-regulatesγ-globin expression in HbE/β-thalassemia patients

To identify crucial miRNAs associated with the wide variation in fetal hemoglobin (HbF) levels observed among HbE/β-thalassemia patients, we performed miRNA expression profiling using the miScript miRNA PCR Array (MIHS-121Z, QIAGEN, GmbH, Hilden, Germany). A total of 16 HbE/β-thalassemia patient samples with varying HbF levels were collected and confirmed to have β-globin and HbE mutations by Sanger sequencing (Figure 3B.ii).Once confirmed as HbE/β-thalassemia cases, patients were initially categorized into two groups, high HbF and normal HbF, based on their γ-globin expression and HPLC profiles. Comparative analysis confirmed that individuals in the high HbF group exhibited significantly elevated HbF levels compared to those in the normal HbF group (Figure 3B.ii). To further investigate the physiological relevance of HbF, we correlated HbF levels with disease severity. Pearson correlation analysis revealed a significant inverse relationship between HbF levels and disease severity, as assessed using the Mahidol scoring system (Figure 3B.i). The miRNA profiling revealed that out of 84 unique miRNAs, 48 were differentially expressed between high and normal HbF groups-31 up-regulated and 17 down-regulated (Table S3). To refine potential candidate miRNAs involved in γ-globin regulation, we conducted parallel miRNA profiling in hydroxyurea (HU)-treated K562 cells, where 45 miRNAs (38 up-regulated, 7 down-regulated) were differentially expressed under high HbF conditions(Table S4). Overlapping miRNAs identified in both patient samples and HU-treated K562 cells which includes miR-148b, miR-320, Let-7b, etc. were shortlisted for subsequent lncRNA–miRNA sponge network analysis.

**Figure 3.**
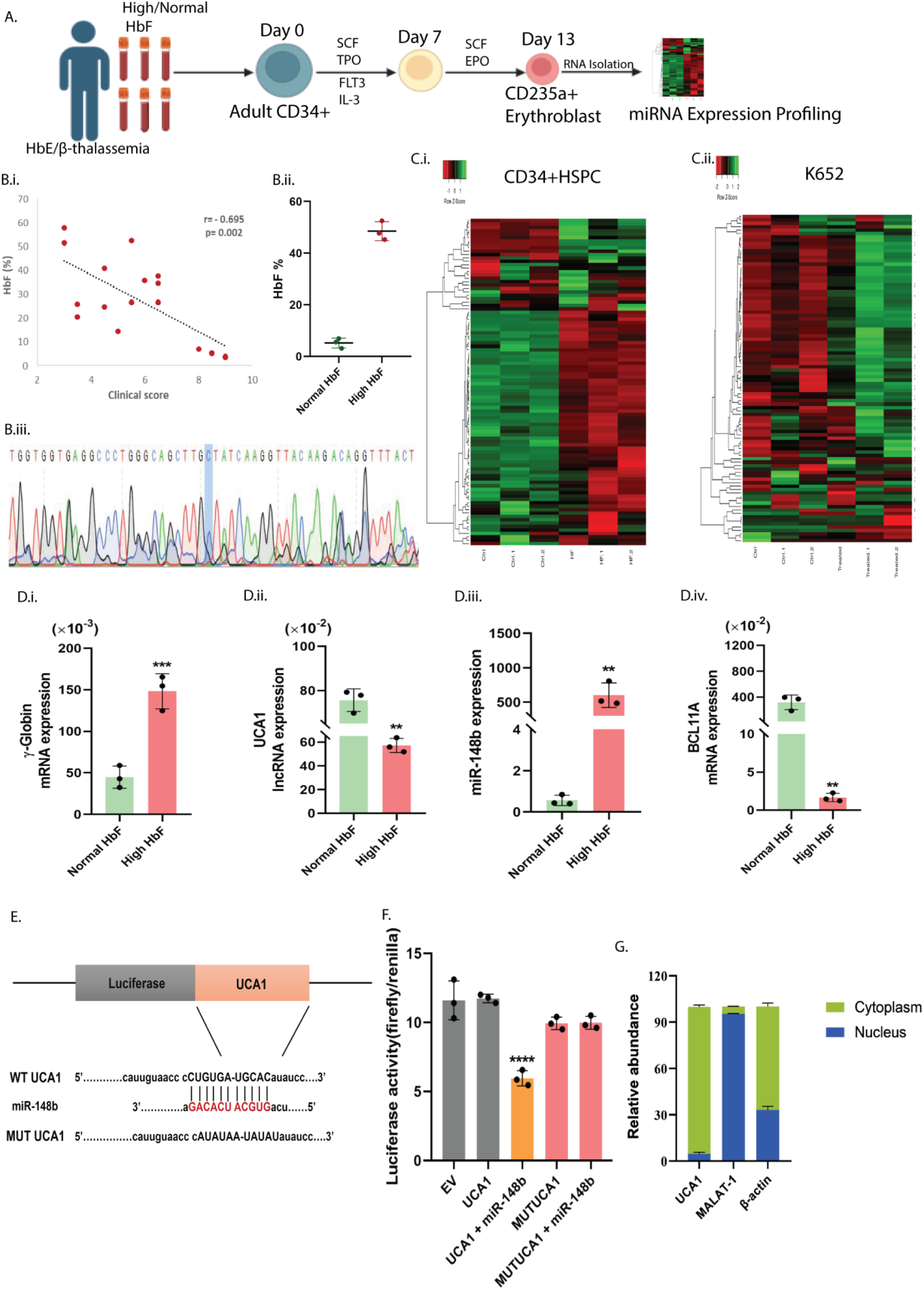
Integrated analysis of miRNA–lncRNA interactions and clinical correlations in CD34⁺ HSPCs collected from HbE/β-thalassemia patients. (A.) Schematic diagram showing miRNA expression profiling from adult CD34+ HSPCs obtained from HbE/β-thalassemia patients. (B.i.) The data is showing associations between HbF levels and clinical score of HbE/β-thalassemia patients. The Pearson’s correlation coefficients were utilized to evaluate the associations of HbF levels with disease severity which is represented as clinical score. The Pearson correlation coefficient (r) and p-value are shown. (B.ii.) Comparison of HbF levels between two groups, patients with normal HbF (<8%) and patients with high HbF (>40%) are shown. (B.iii.) Representative Chromatogram of DNA sequences denoting HBB mutation in patient sample. Direct Sequencing analysis revealed the presence of IVS-1-5(G>C) mutation in Intron 1 of β-globin gene. The blue box marked the position of the substitution in β-globin gene. (C.i-ii). Heat map showing differentially expressed miRNAs in (i) patients(high HbF and normal HbF) derived CD34+ HPSCs cells, and in (ii) K562 cells post HU-treatment. (D.i-iv). Figure showing expression of (i) γ-globin, (ii) UCA1, (iii) miRNA, and (iv) BCL11A in high HbF and normal HbF conditions (patient CD34+ HSPCs). (E.) Figure showing sequence of UCA1 with putative binding site of miR-148b. MUTUCA1 refers to a mutant variant of UCA1 in which the miR-148b binding sites have been disrupted. (F.) Figure showing luciferase assay to confirm direct interaction between UCA1 and miR-148b. (G.) Figure showing relative abundance of UCA1 in cytoplasmic and nuclear fractions of K562 cells. Data are representative of three independent biological replicates and graphical data are represented as Mean ± SD. * p <0.05, ** p <0.01, **** p<0.0001 by unpaired two-tailed t-test.

UCA1 has previously been reported to act as sponges to bind miRNAs and therefore regulate their functions in cancer[30]. Therefore, we hypothesized that UCA1 may have binding sites for differentially expressed miRNAs identified in our study. In order to construct lncRNA-miRNA networks, we assessed miRNA expressions in erythroid cells and found that miR-148b showed opposite expression pattern compared to that of UCA1. Subsequent bioinformatics analysis revealed putative binding sites for miR-148b in UCA1. RT-qPCR analysis confirmed the opposite expression patterns of UCA1 and miR-148b in HbE/β-thalassemia patients with high HbF when compared with patients with normal HbF (Figure 3D). In addition, association between the UCA1-lncRNA/miR-148b/BCL11A axis and HbF levels in HbE/β-thalassemia patients revealed that UCA1 and BCL11A exhibit negative correlations, while miR-148b shows a positive correlation with elevated HbF levels (Figure S2).Since sub-cellular localisation can provides insight regarding the function of lncRNA, we performed cell fractionation study to assess the distribution of UCA1 in maturing erythroid cells. Our results showed that UCA1 expressed predominantly in cytoplasm of adult erythroid cells, supporting the notion that UCA1 may act as miR-148b sponge in adult erythroid cells (Figure 3G).

### UCA1 directly binds with miR-148b

Long non-coding RNAs (lncRNAs) can function as competing endogenous RNAs (ceRNAs) by sequestering shared microRNAs (miRNAs) through complementary base-pairing, thereby regulating gene expression post-transcriptionally. To investigate whether UCA1 can act as a molecular sponge for miR-148b, we first performed bioinformatics analyses, which predicted the presence of putative miR-148b binding sites within the UCA1 transcript. To experimentally validate this interaction, a fragment of UCA1 containing the predicted miR-148b binding sites was cloned downstream of the firefly luciferase gene (luc2) in the pmiRGLO dual-luciferase reporter vector. This vector also contains a constitutively expressed Renilla luciferase gene (Rluc) for normalization of transfection efficiency. The UCA1-pmiRGLO plasmid was co-transfected into cells along with synthetic miR-148b mimics, and luciferase activity was measured to assess direct interactions. Overexpression of miR-148b led to a dose-dependent decrease in firefly luciferase activity, indicating that miR-148b binds to the UCA1 reporter construct and represses gene expression. To confirm that this repression was specifically due to the predicted miR-148b binding sites, we generated a mutant construct in which the seed region complementary to miR-148b was altered. Co-transfection of this mutant construct with miR-148b mimics showed no significant reduction in luciferase activity compared to the empty vector control (Figure 3F), thereby confirming the specificity of the interaction.Collectively, these findings demonstrate that UCA1 directly interacts with miR-148b through complementary base-pairing, supporting a ceRNA-based regulatory mechanism involving the UCA1–miR-148b axis.

### UCA1 negatively regulates fetal haemoglobin expression by sponging miR-148b in erythroid cells

To investigate whether UCA1 acts as an endogenous sponge for miR-148b and thereby suppresses miR-148b-mediated induction of fetal haemoglobin (HbF), we performed a gain-of-function study by overexpressing full-length UCA1 cloned into a pcDNA3.1-mCherry vector, downstream of the 3′UTR of mCherry. Overexpression was confirmed by RT–qPCR (Figure 4B.ii), and stable expression of the UCA1–mCherry fusion transcript was visualised by confocal microscopy (Figure 4A). Compared to control cells, UCA1-overexpressing K562 cells exhibited a significant reduction in miR-148b and γ-globin transcript levels, indicating that UCA1 suppresses γ-globin expression by antagonising miR-148b (Figure 4B.v). Immunoblot analysis corroborated the transcriptional data, showing reduced γ-globin protein levels in UCA1-overexpressing cells, even upon miR-148b mimic transfection. This suppression recapitulated the effect of miR-148b antagomiR co-transfection (Figure 4C). To complement the gain-of-function data, we performed a loss-of-function study using siRNA-mediated knockdown of UCA1 (si-UCA1) in K562 cells. Knockdown of UCA1 led to elevated γ-globin expression, as assessed by both RT–qPCR and Immunoblot analyses, further supporting the role of UCA1 as a negative regulator of HbF through sequestration of miR-148b (Figure 4D-E).To further evaluate whether the effects on γ-globin expression were due to sponging between overexpressed UCA1 and miR-148b, RNA-RNA pull down assays were performed on extract of K562 cells overexpressing UCA1 using biotinylated miRNA mimics (Figure 4F.i). The RT-qPCR results showed that UCA1 expression was preferentially enriched in the biotinylated miR-148b compared to the non-targeting mimics. These results suggest UCA1 enrichments in biotinylated miR-148b, were likely due to direct association of UCA1 with miR-148b (Figure 4F.ii)

**Figure 4.**
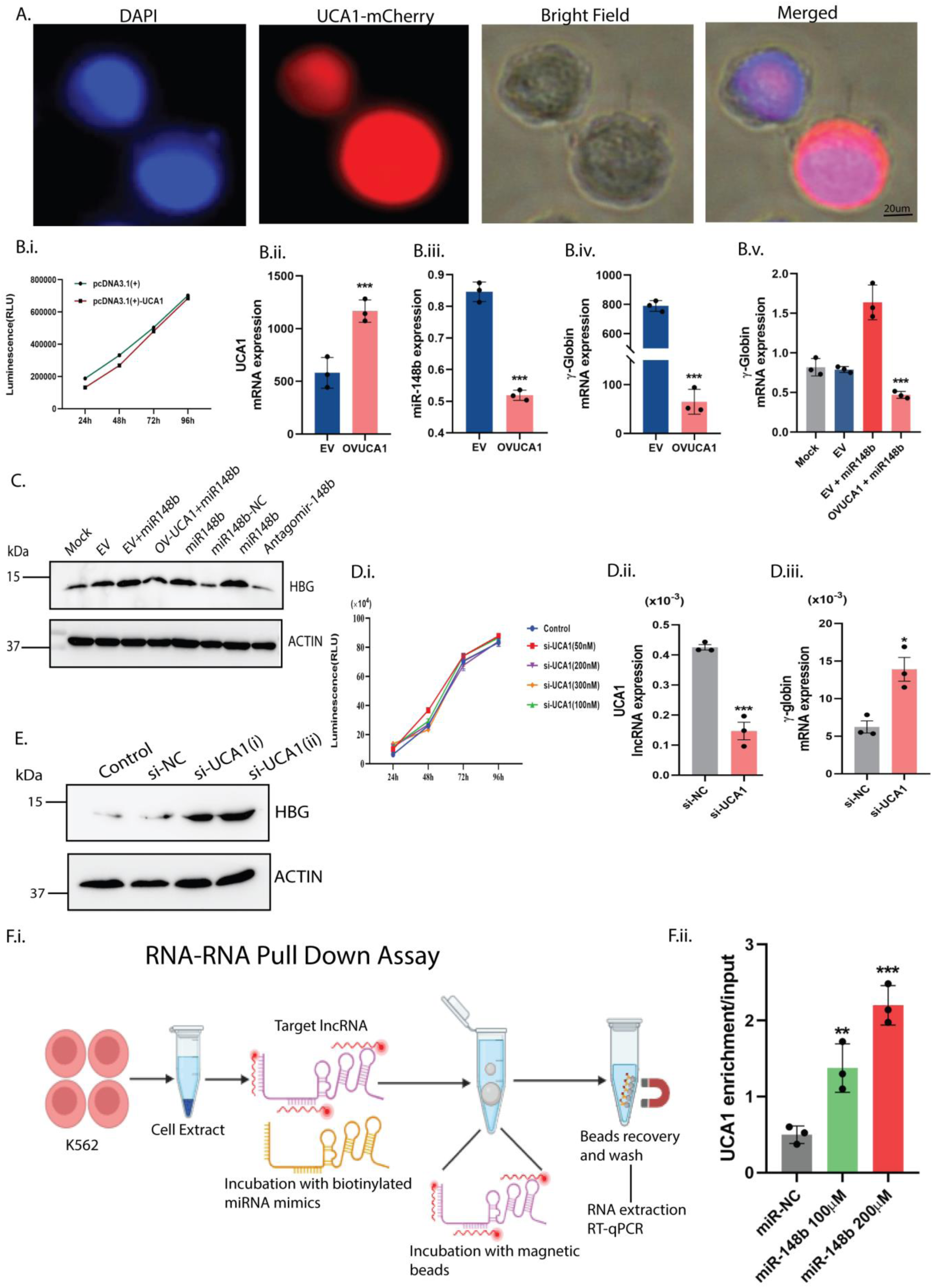
UCA1 over-expression/knock-down modulates γ-globin expression via sequestering miR-148b. (A.) Confocal image showing localisation of UCA1-mCherry (red) around cytoplasm with DAPI, representing nucleus in K562 cells. (B.i.) Cell Titer®-Glo assay results showing proliferation of the K562 cells at different time points post transfection with empty vector, pcDNA3.1(+) and UCA1-cloned vector, UCA1-pcDNA3.1(+). (B.ii-v) RT-qPCR analysis showing expression levels of (ii) UCA1, (iii) miR-148b, and (iv) γ-globin following UCA1 overexpression in K562 cells. (B.v) RT-qPCR analysis showing γ-globin expression in UCA1-overexpressing cells following transfection with miR-148b mimic. (C.i) Immunoblots are showing γ-globin and β-actin protein levels in miR-148b-transfected and UCA1-overexpressing K562 cells. (D.i) CellTiter®-Glo assay showing proliferation of K562 cells at different time points post-transfection with increasing concentrations (50–300 nM) of siRNA targeting UCA1 lncRNA. (D.ii-iii) RT-qPCR analysis showing expression levels of (ii) UCA1 and (iii) γ-globin following si-UCA1 or negative control (si-NC) transfections. (E.) Immunoblot analysis showing expression of γ-globin and β-actin levels in K562 cells post-siRNA transfection. (F.i) Schematic diagram showing steps of RNA-RNA pull down assay which was used to confirm direct association between UCA1 and miR-148b. (F.ii) RT-qPCR analysis showing UCA1 enrichments in biotinylated miR-148b when compare with biotinylated scramble(miR-NC) in dose-dependent manner. Data are representative of three independent biological replicates and graphical data are represented as Mean ± SD. * p <0.05, ** p <0.01, **** p<0.0001 by unpaired two-tailed t-test.

### UCA1/miR-148b modulates γ-globin expression without perturbing Cell Cycle progression or Apoptosis

To determine whether miR-148b could upregulate γ-globin expression independently of UCA1, we ectopically overexpressed miR-148b in K562 cells and compared γ-globin levels across three experimental conditions: scramble control, UCA1 knockdown, and miR-148b overexpression.

At first, we evaluated the effects of miRNA mimics on cell viability and proliferation. No significant differences were observed between cells transfected with miR-148b mimics, non-targeting mimics, or untreated controls in terms of viability or proliferation (Figure 5B). Similarly, knockdown of UCA1 using siRNA (si-UCA1) did not affect cell viability or proliferation (Figure 4D.i).To assess potential effects on cell cycle progression, we performed flow cytometric analysis of total DNA content of the transfected cells. The cell cycle profiles of both miR-148b mimic- and si-UCA1-transfected cells were comparable to those of controls, with the majority of cells in the G2/M phase (Figure 5D), suggesting that neither miR-148b overexpression nor UCA1 depletion significantly impacts cell cycle dynamics. Moreover, apoptosis levels remained unaltered in both conditions, indicating that these manipulations do not trigger cell death pathways. We next assessed γ-globin expression following transfection. Immunoblot analysis revealed a significant increase in γ-globin levels in cells transfected with either miR-148b mimic or si-UCA1 compared to non-targeting controls (Figure 5E). This up-regulation was further corroborated by intracellular fetal hemoglobin (F-cells) staining and flow cytometry, which showed a marked increase in the percentage of F-cells in both conditions (Figure 5F). Collectively, these findings demonstrate that both miR-148b overexpression and UCA1 knockdown significantly enhance HbF expression in erythroid cells, without perturbing cell cycle progression or inducing apoptosis, underscoring their regulatory role in γ-globin gene expression.

**Figure 5.**
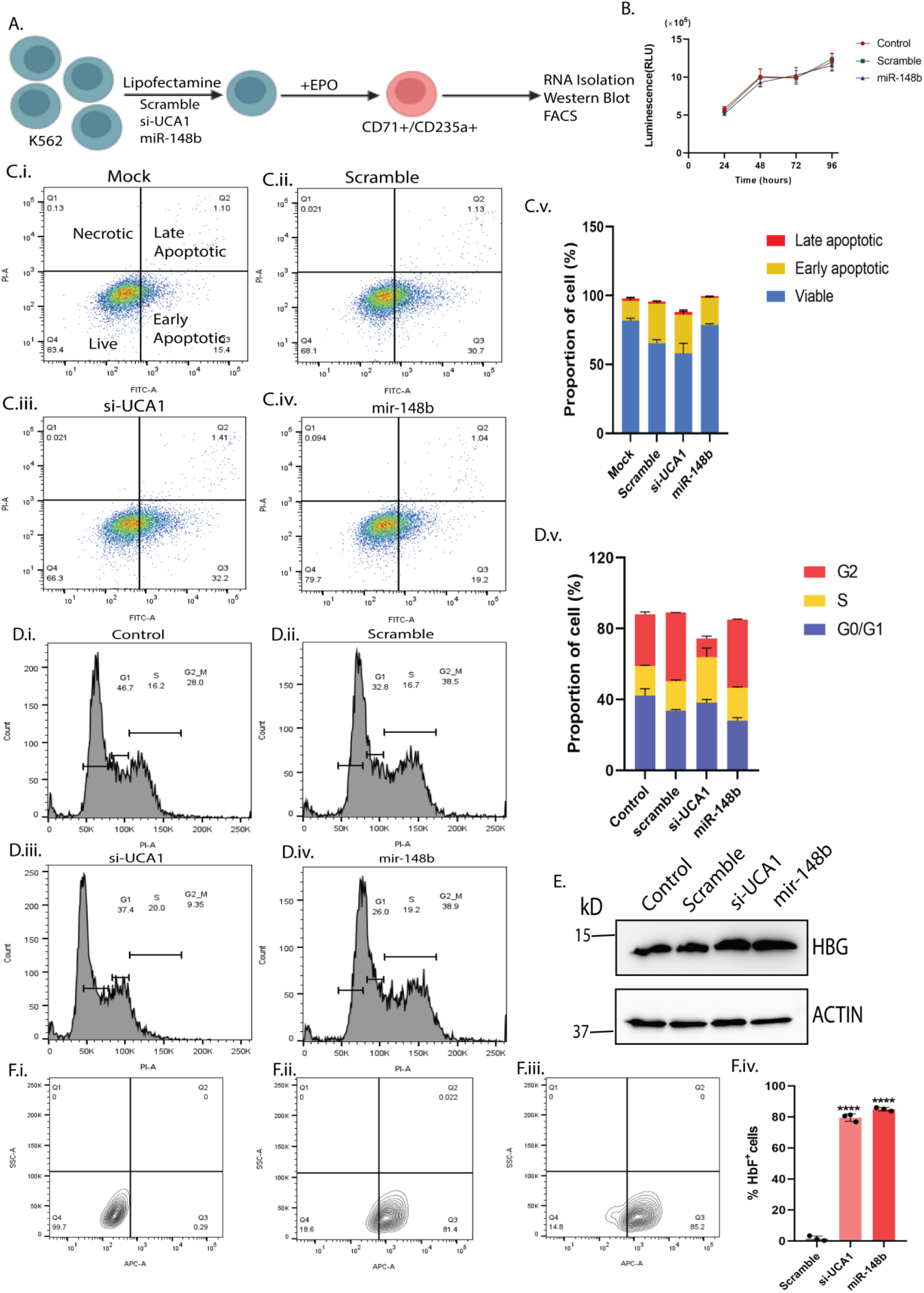
Figure showing functional effects of UCA1 knockdown and miR-148b overexpression on apoptosis, cell cycle, and γ-globin induction in K562 cells. (A.) Schematic diagram showing transfections of scramble, miR-148b mimics and si-UCA1 in K562 cells. (B.) Cell Titer®-Glo assay results showing proliferation of the cells at different time points post transfection with scramble control and mir-148b mimics. (C.i-v) FACS plots showing percentage of apoptotic cells in (i) mock, (ii) Scramble control (iii) si-UCA1, and (iv) miR-148b mimics. (v) Bar diagram showing different proportions of viable cells in mock, scramble, miR-148b mimics and si-UCA1 transfected cells. (D.i-v) Flow cytometry analysis of cell cycle progression post transfection. Histograms showing different proportions of cell cycle phases (G1, S and G2M) in (i) Mock, (ii) Scramble control, (iii) si-UCA1 and (iv) miR-148b mimics, respectively. (v) Bar graphs showing different proportions of cell cycle phases post scramble, si-UCA1 and miR-148b transfection. (E.) Immunoblot analysis showing expression of γ-globin and β-actin levels in K562 cells post scramble control, si-UCA1 and miR-148b transfection. (F.i-iii) Quantification of percentage of HbF positive cells (F-cells) assessed by flow cytometry analysis in (i) Scramble, (ii) si-UCA1 and (iii) miR-148b treated K562 cells. (F.iv) Bar graphs showing percentage of F-cells post miRNA-transfection. Data are representative of three independent biological replicates and graphical data are represented as Mean ± SD. * p <0.05, ** p <0.01, **** p<0.0001 by one-way ANOVA.

### BCL11A was confirmed as a direct downstream target of miR-148b through a luciferase reporter assay

To uncover specific gene targets of miR-148b involved in γ-globin regulation, we employed the multiMiR R package and database; a comprehensive repository integrating predicted and validated miRNA-target interactions from 14 different sources. This analysis identified a total of 207 candidate genes containing putative miR-148b binding sites within their 3′ untranslated regions (3′UTRs). Given that miR-148b expression is elevated under high HbF conditions, we hypothesized that its target genes might exhibit stage-specific expression profiles that are inversely correlated with miR-148b levels. To explore this, we intersected the list of 207 predicted targets with genes that were previously found to be differentially expressed in our transcriptome analysis of high-HbF versus low-HbF conditions. This integrative approach narrowed down the candidate list to five genes-BCL11A, ZBTB7A, SOX6, TAL1, and MYB-all of which were significantly deregulated in high-HbF conditions and contained miR-148b binding sites in their 3′UTRs. Among these, BCL11A and ZBTB7A are well-established transcriptional repressors of γ-globin, known to directly bind to the γ-globin promoter. Therefore, these two candidates were prioritized for experimental validation. To assess whether miR-148b can regulate the expression of BCL11A and ZBTB7A via direct interaction with their 3′UTRs, we performed luciferase reporter assays in HEK293 cells. Fragments of the 3′UTRs containing the predicted miR-148b seed-matching regions were cloned downstream of the firefly luciferase (luc2) gene in the pmiRGLO vector, and co-transfected with either miR-148b mimics or non-targeting control mimics. As shown in Figure 6C.i, co-transfection with miR-148b significantly suppressed luciferase activity from the BCL11A-3′UTR reporter compared to the scramble control, indicating direct targeting. Furthermore, deletion of the miR-148b seed site within the BCL11A 3′UTR abrogated this repression, confirming the specificity of the interaction. In contrast, a similar luciferase assay using the ZBTB7A-3′UTR reporter showed no significant reduction in luciferase activity upon miR-148b mimic transfection (Figure 6C.ii), suggesting that despite the predicted binding site, ZBTB7A is not a direct functional target of miR-148b in this context.

**Figure 6.**
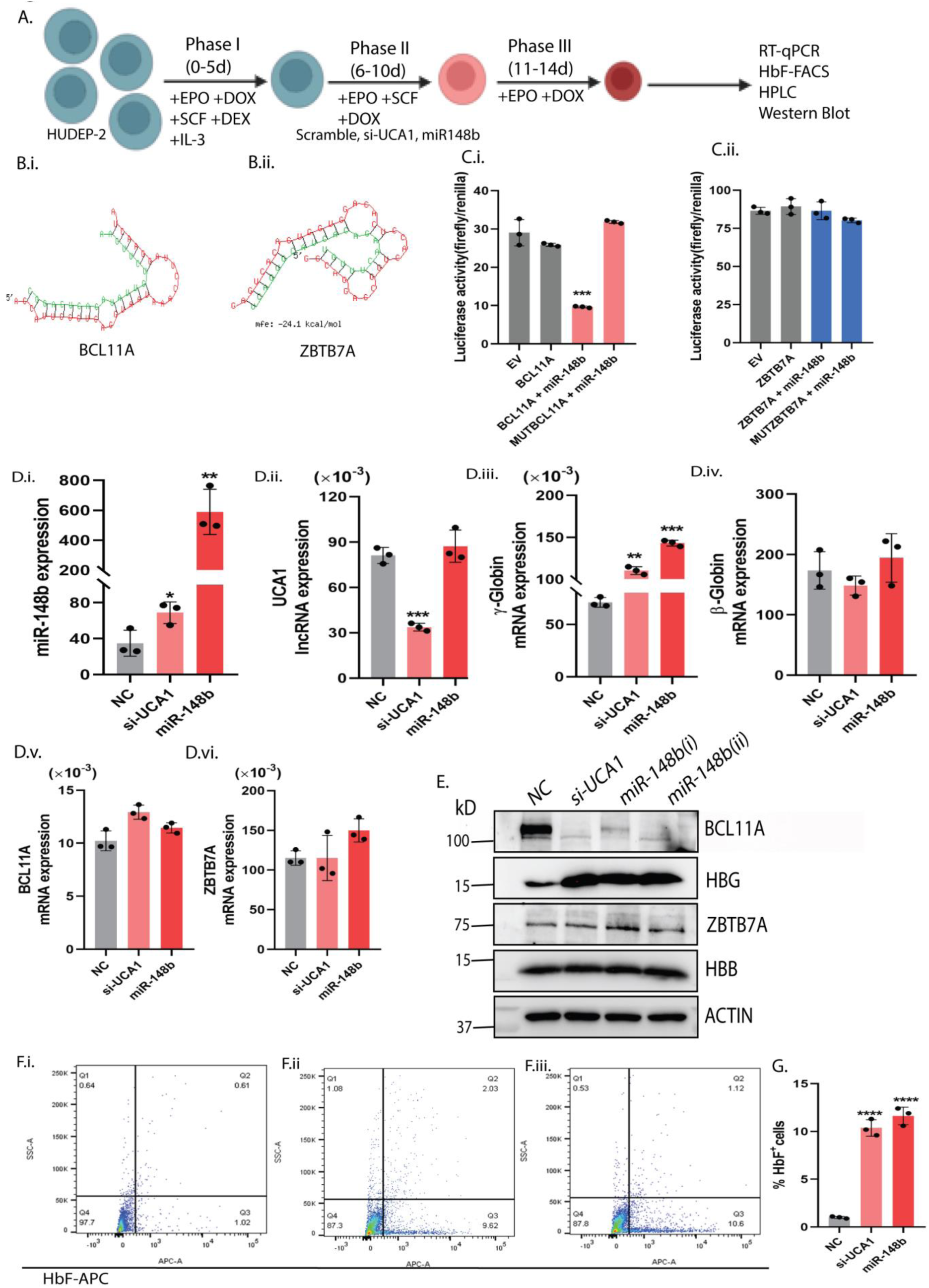
UCA1 knockdown or miR-148b overexpression promotes HbF induction in HUDEP-2 cells via suppression of BCL11A. (A.) Schematic diagram showing HUDEP-2 cells proliferation and differentiation in specific media. (B.і-іі) RNAhybrid results showing base pairing between miR-148b seed region and 3’UTR of target genes (BCL11A and ZBTB7A). (C.i-ii) Figure is showing luciferase assay results which confirm the direct interactions between miR-148b and BCL11A 3’UTR. (ii) Figure showing luciferase assay results which confirm no direct interactions between miR-148b and ZBTB7A 3’UTR. (D.i-vi) RT-qPCR analysis of (i) miR-148b, (ii) UCA1, (iii) γ-globin, (iv)β-globin, (v)BCL11A and (vi)ZBTB7A expression in HUDEP-2 cells post miR-148b and si-UCA1 transfections. Results are normalized to endogenous β-actin expression. (E.) Immunoblot analysis showing expression of BCL11A, HBG, ZBTB7A, HBB and Beta-actin in UCA1 depleted (si-UCA1) and miR-148b overexpressed HUDEP-2 cells. Signals from Immunoblots are normalized to endogenous β-actin expression. (F.i-iii) Representative flow cytometry plots showing anti-HbF-staining in (i) Negative control, (ii) si-UCA1, and (iii) miR-148b-transfected HUDEP-2 cells. (G) Bar plots showing percentage of HBG-containing F-cells measured by flow cytometry analysis in (i) Negative control, (ii) si-UCA1, and (iii) miR-148b-transfected HUDEP-2 cells. Data are representative of three independent biological replicates and graphical data are represented as Mean ± SD. * p <0.05, ** p <0.01, **** p<0.0001 by one-way ANOVA.

Together, these findings demonstrate that BCL11A is a bona fide target of miR-148b, which can directly suppress its expression through interaction with the 3′UTR, thereby potentially contributing to γ-globin de-repression. No such regulatory effect was observed for ZBTB7A, highlighting the specificity of miR-148b-mediated post-transcriptional regulation

### UCA1 depletion/miR-148b overexpression elevates fetal hemoglobin expression via BCL11A repression in HUDEP-2 cells

To demonstrate that miR-148b directly represses BCL11A expression, we ectopically overexpressed miR-148b in adult-type erythroid HUDEP-2 cells. Successful overexpression was first confirmed by RT-qPCR analysis (Figure 6D.i). Transfection with miR-148b, but not a non-targeting control, significantly up-regulated γ-globin expression in HUDEP-2 cells (Figure 6D.iii). As expected, no changes were observed in the expression levels of β-globin or ZBTB7A (Figure 6D.iv-vi). Immunoblot analysis further supported these findings, showing a dose-dependent increase in γ-globin expression following miR-148b transfection (Figure 6E). Interestingly, the levels of BCL11A primary transcript remained unchanged upon miR-148b overexpression compared to the negative control. However, BCL11A protein expression was significantly reduced in miR-148b–transfected cells, indicating post-transcriptional regulation of BCL11A by miR-148b (Figure 6E). Given that miR-148b modulates BCL11A post-transcriptionally, potentially leading to HBG induction, we next assessed fetal hemoglobin (HbF) levels by F-cell staining. The flow cytometry analysis confirmed a significant increase in HbF levels in cells overexpressing miR-148b.

As we had previously established that the long non-coding RNA UCA1 acts as a molecular sponge for miR-148b, we next sought to determine whether knockdown of UCA1 (si-UCA1) could recapitulate the effects of miR-148b overexpression in HUDEP-2 cells. RT-qPCR confirmed efficient silencing of UCA1 (Figure 6D.ii). Remarkably, si-UCA1 treatment alone resulted in significant up-regulation of γ-globin expression, with no observable changes in β-globin or ZBTB7A, phenocopying the effect of miR-148b (Figure 6D.iii-vi). Immunoblot analysis further confirmed elevated γ-globin protein levels in si-UCA1–treated cells, along with a substantial reduction in BCL11A protein, while transcript levels remained unaffected, again indicating post-transcriptional regulation (Figure 6E). Consistently, F-cell staining demonstrated a significant increase in HbF levels in response to UCA1 knockdown (Figure 6F.i-iii). No overt changes were observed in cell viability or erythroid maturation upon miR-148b overexpression and/or UCA1 depletion, as assessed by cell surface phenotyping (CD235a and CD71) or CellTitre GLO viability assay on day 14 of differentiation (Supplemental figure S4-S5).

Together, these findings suggest that both miR-148b overexpression and UCA1 knockdown independently induce HbF production by down-regulating BCL11A expression at the post-transcriptional level. These results highlight a convergent regulatory axis in which UCA1 functions upstream of miR-148b to modulate fetal hemoglobin induction in adult erythroid cells.

### UCA1 depletion/miR-148b overexpression reactivates fetal hemoglobin in adult primary human erythroblasts

To further validate our findings in a physiologically relevant system, we extended our mechanistic studies to primary human erythroblasts differentiated ex-vivo from CD34⁺ hematopoietic stem and progenitor cells (HSPCs) isolated from healthy adult donors. These primary cells provide a robust model for studying erythroid biology and hemoglobin switching in a setting that closely mirrors in vivo erythropoiesis. CD34⁺ HSPCs were cultured under standard erythroid differentiation conditions and transfected with a non-targeting scramble control, synthetic miR-148b mimics, or siRNA targeting UCA1 (si-UCA1). Cell viability and erythroid differentiation status were assessed at day 6 of differentiation using flow cytometry analysis of lineage-specific markers CD71 and CD235a, along with cytological examination following Wright-Giemsa staining (Figure 7B and Supplemental Figure S7). Importantly, neither miR-148b overexpression nor UCA1 knockdown had a significant effect on cell viability or erythroid maturation, suggesting that these interventions do not impair normal erythroid development. These findings are crucial in establishing the therapeutic feasibility of manipulating the miR-148b–UCA1 axis without deleterious effects on erythropoiesis.

**Figure 7.**
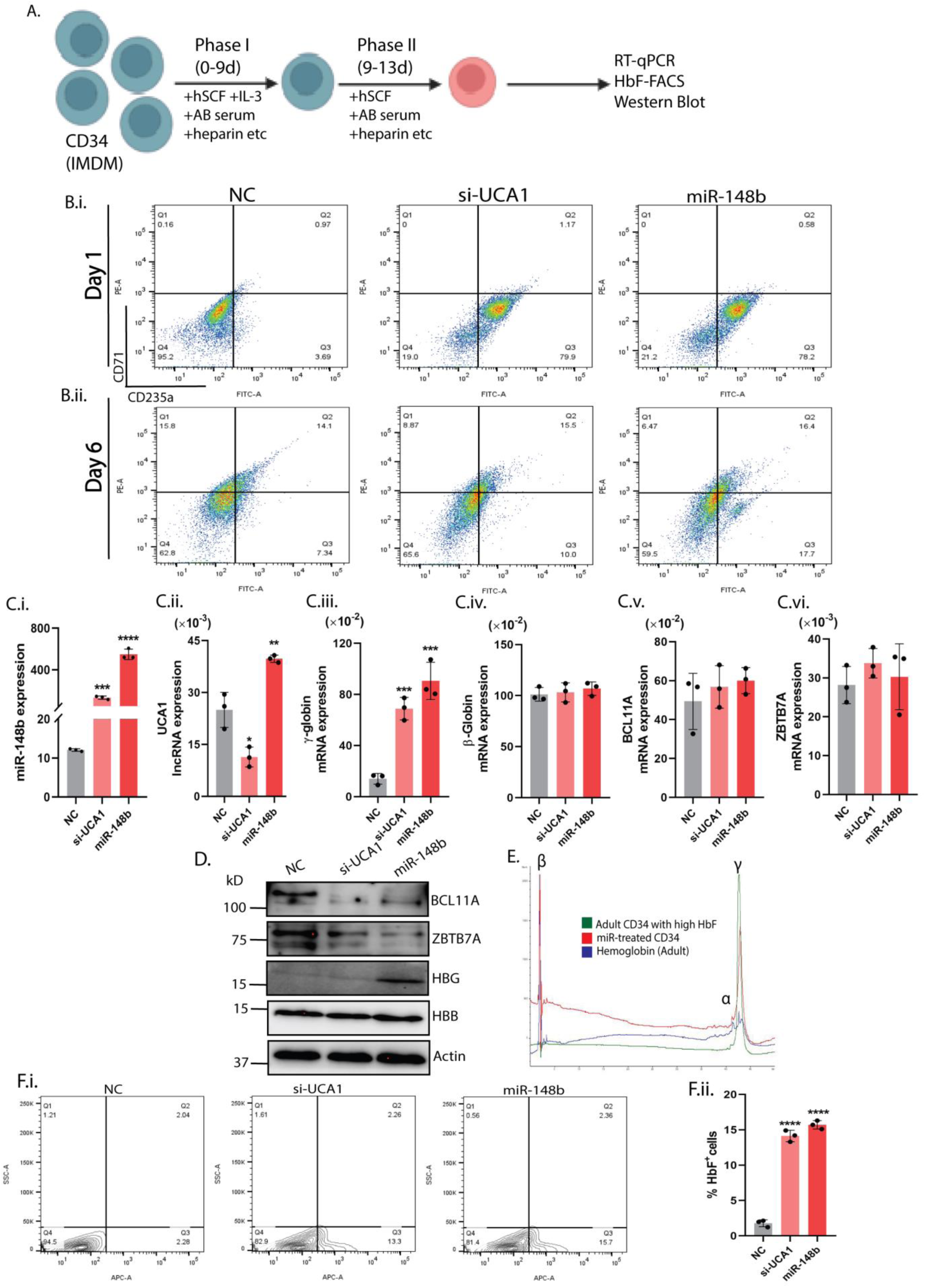
UCA1 knockdown or miR-148b overexpression promote HbF induction in adult primary erythroblasts. A Schematic diagram showing *in-vitro* expansion and differentiation of CD34+ HSPCs derived from healthy adults. (B.i-ii). Representative flow cytometry of CD235a and CD71 double staining of primary erythroblasts at day 1 and 6 of differentiation. (C.i-vi). RT-qPCR analysis of (i) miR-148b, (ii) UCA1, (iii) γ-globin, (iv) β-globin, (v) BCL11A and C.vi. ZBTB7A expression in adult human primary erythroblasts post miR-148b and si-UCA1 transfection. Results are normalized to endogenous control, β-actin expression and shown as mean ± SD. (D.) Representative Immunoblots with indicated antibodies of extracts from primary human erythroblasts transfected with scramble, si-UCA1 and miR-148b mimics. Signals from Immunoblots are normalized to endogenous control, β-actin expression. Results are represented in three biological replicates. (E.) HbF induction in primary erythroblasts with miR-148b transfection measured by HPLC analysis. (F.i.) Representative flow cytometry plots showing anti-HbF-staining in (i) Negative control, (ii) si-UCA1, and (iii) miR-148b-transfected primary erythroblasts derived from CD34+ HSPCs. (G) Bar plots showing percentage of HBG-containing F-cells measured by flow cytometry analysis in (i) Negative control, (ii) si-UCA1, and (iii) miR-148b-transfected primary erythroblasts derived from CD34+ HSPCs. Results are shown as mean ± SD. Data are representative of three independent biological replicates and graphical data are represented as Mean ± SD. * p <0.05, ** p <0.01, **** p<0.0001 by one-way ANOVA.

We next investigated whether these interventions could induce fetal hemoglobin (HbF) expression in primary erythroblasts. RT-qPCR analysis revealed a substantial up-regulation of HBG1/2 mRNA upon transfection with miR-148b mimics and/or si-UCA1, relative to scramble controls (Figure 7C). This increase in γ-globin transcription was corroborated at the protein level by Immunoblot analysis, which demonstrated elevated γ-globin levels in treated cells. Notably, expression of HBB (β-globin) and ZBTB7A (a known HbF repressor) remained largely unchanged, indicating that miR-148b/UCA1 perturbation selectively affects γ-globin regulation without broadly altering the expression of other globin genes or transcriptional repressors (Figure 7D). Flow cytometric analysis using an anti-HbF antibody further revealed a significant increase in the proportion of HbF-positive cells (F-cells) in both miR-148b and si-UCA1 treated groups (Figure 7F.i-ii). Strikingly, HPLC-based quantification demonstrated that miR-148b overexpression alone elevated the proportions of γ-globin to an extent comparable to that observed in individuals with naturally high HbF levels, highlighting its strong therapeutic potential(Figure 7E). To elucidate the mechanistic basis of γ-globin reactivation, we assessed the impact of these interventions on BCL11A, a master repressor of HbF. Consistent with our earlier findings in HUDEP-2 cells, BCL11A protein levels were markedly reduced in miR-148b- and si-UCA1-transfected primary erythroblasts, whereas levels of the BCL11A primary transcript remained unaltered, as determined by RT-qPCR (Figure 7C.v). This observation supports a model in which the miR-148b–UCA1 axis exerts post-transcriptional silencing of BCL11A, leading to de-repression of the γ-globin gene.

Collectively, these results reinforce the role of the miR-148b–UCA1–BCL11A regulatory axis in fetal hemoglobin induction within primary CD34⁺-derived erythroid cells and underscore the translational potential of RNA-based approaches for HbF reactivation in β-hemoglobinopathies.

## DISCUSSION

Human fetal hemoglobin (HbF) is developmentally silenced after birth, and reactivation of fetal hemoglobin genes in adult stage could ameliorate severe symptoms associated with β-hemoglobin disorders, such as SCD or β-thalassemia[31]. During development, γ-globin genes (HBG) are tightly regulated by different cis-regulatory and trans-acting factors, including lineage-specific transcription factors and co-factors such as BCL11A, ZBTB7A, GATA1, KLF1, MYB, etc. Over the past decades, BCL11A and ZBTB7A have been identified to be major repressors that play a crucial role in the fetal-to-adult hemoglobin switching, and depletion of one of these repressors can independently de-repress γ-globin expression in adult erythroid cells[32].Accumulating evidence, particularly from genome editing and transcriptomic analyses, indicates that erythroid-specific attenuation of BCL11A levels can induce sustained HbF expression, offering durable therapeutic benefit in preclinical and clinical settings[33].While transcriptional repressors such as BCL11A and ZBTB7A are well-established mediators of γ-globin silencing, the upstream regulatory networks that fine-tune their expression remain incompletely understood. Here, we identify a post-transcriptional regulatory axis involving the long non-coding RNA UCA1 and miR-148b, which modulates BCL11A expression and contributes to the γ-globin gene silencing in adult erythroid cells. Our study demonstrates that UCA1, a developmentally regulated and abundantly expressed lncRNA in erythroid cells, functions as a molecular sponge for miR-148b. By sequestering miR-148b, UCA1 indirectly stabilizes BCL11A transcripts, thereby reinforcing γ-globin silencing. The integrative analysis of lncRNA and miRNA expression profiles in primary CD34+ hematopoietic cells and erythroid cell lines further supports a co-regulated ceRNA network with UCA1 and miR-148b at its core. The physiological relevance of this interaction is supported by loss- and gain-of-function experiments in erythroid model systems (CD34+HSPCs and HUDEP-2), where UCA1 depletion or miR-148b overexpression led to significant induction of γ-globin. These findings highlight a previously unappreciated layer of post-transcriptional control in globin gene regulation, expanding the mechanistic understanding of how HbF expression is fine-tuned during erythroid maturation. Studies have shown that cellular miRNA abundance could be fine-tuned for regulatory effects using ectopically expressed artificial transcripts (miRNA sponges) containing tandem repeats of miRNA binding sites[35][36]. In addition, several naturally occurring mRNAs, transcribed pseudogenes, and lncRNAs have already been identified as miRNA decoys, regulating various cellular processes by sequestering miRNAs, and thereby protecting their target mRNAs from repression[37][38].

While prior studies have reported the utility of artificial miRNA sponges and pseudogene-derived decoys in modulating miRNA activity, our data identify UCA1 as a naturally occurring lncRNA sponge that modulates a clinically relevant target in the context of hemoglobin switching. Importantly, the predicted binding sites of miR-148b on UCA1 transcript lack strong evolutionary conservation, yet our luciferase assays and cross-linking affinity purification experiments provide evidence of direct interactions, reinforcing the notion that non-canonical and non-conserved miRNA recognition sites can retain functionality in a species- and cell-type-specific manner[39][40]. Our results contribute to both the growing list of functional non-canonical sites and provide evidence for the indispensable role of non-conserved seed sites in non-coding RNA-mediated sponge activity. In addition to its known role in stabilizing ALAS2 mRNA during heme biosynthesis via PTBP1 recruitment[34], our data unveil a distinct function of UCA1 in regulating miRNA activity. This context-dependent functional versatility underscores the broader regulatory potential of lncRNAs as integrators of both transcriptional and post-transcriptional mechanisms.

In summary, this study identifies UCA1 as a previously unrecognized regulator of γ-globin (HBG1/2) gene, functioning through miR-148b sequestration to maintain BCL11A expression in adult erythroid cells (Figure 8). These findings elucidate a novel layer of post-transcriptional control within the globin gene regulatory network and open new avenues for RNA-based therapeutic strategies in β-thalassemia and sickle cell disease. By expanding our understanding of non-coding RNA-mediated regulation, this work contributes to the growing appreciation of lncRNAs as key players in developmental gene control and disease modulation.

**Figure 8.**
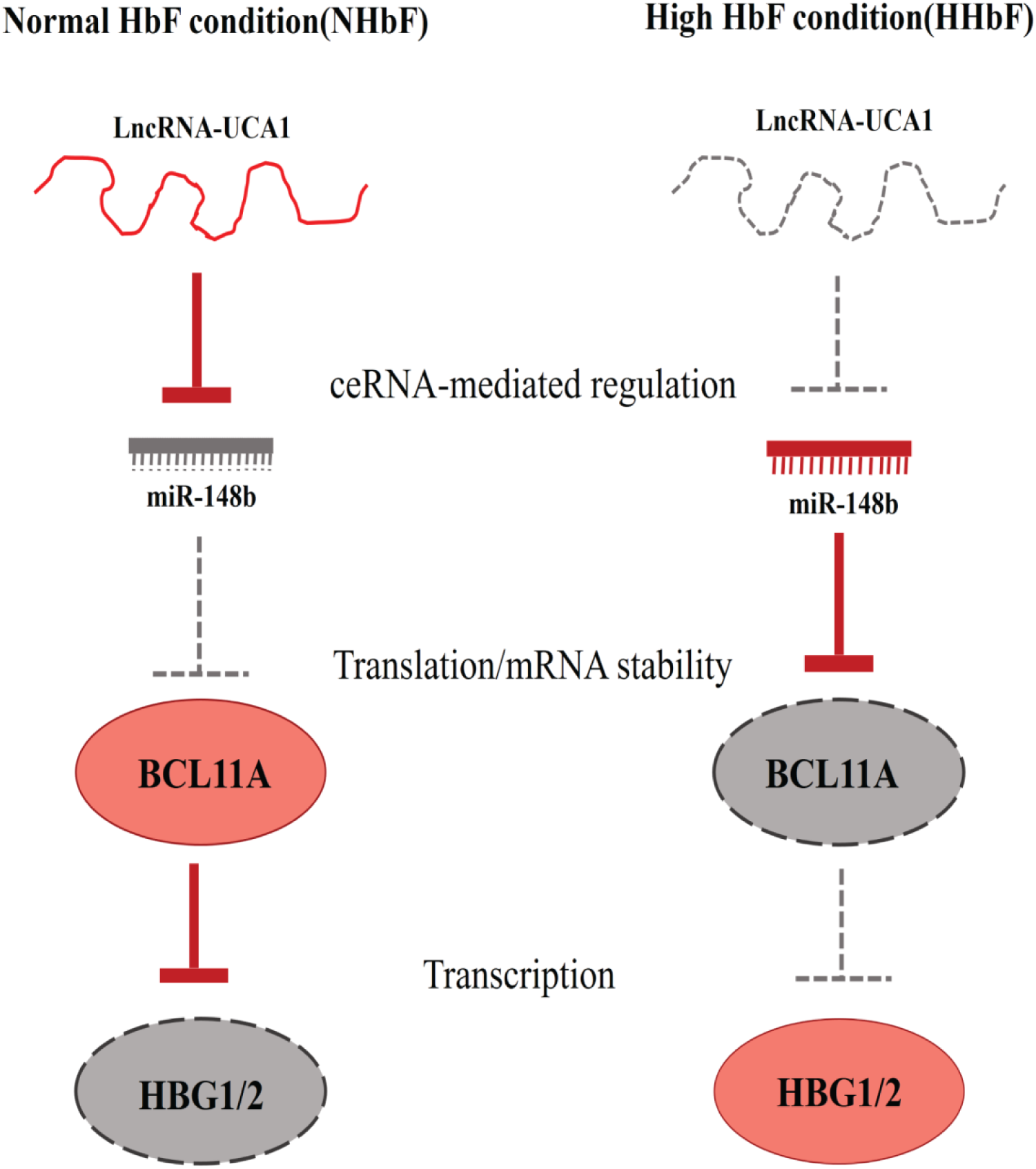
Schematic representation of HbF regulation via the UCA1/miR-148b mediated competing endogenous RNA (ceRNA) network. In normal (low HbF) conditions, the lncRNA UCA1 sequesters miR-148b, preventing it from downregulating BCL11A, a key repressor of HBG (γ-globin). This results in high BCL11A expression and suppression of HBG transcription. In contrast, during elevated HbF conditions, UCA1 is downregulated, leading to increased availability of miR-148b, which represses BCL11A and subsequently derepresses HBG, thereby promoting γ-globin expression.

## MATERIALS AND METHODS

### Study design

Peripheral blood was collected from six unrelated HbE/β-thalassemia patients with varying HbF levels. Primary screening followed by molecular analysis was performed to confirm the disease. All patient samples were categorized either as normal HbF (NHbF: HbF<10%, n=3) and high HbF (HHbF: HbF>40%, n=3) groups. The demographic details and clinical features of the patients were enlisted in the Supplemental information (Table S1). Peripheral blood from another 10 unrelated individuals with HbE/β-thalassemia comprising of normal- and high-HbF groups was used for correlation study. In addition, all patients were assigned clinical scores based on the Mahidol Scoring system to assess the association between HbF levels and disease severity. Written informed consent was taken from each participant and/or their guardians. The study was conducted according to the guidelines provide by the Declaration of Helsinki.

### Cell culture

All cell culture was performed at 37^0^ C with 5% CO_2_ in water-jacketed CO_2_ incubator under sterile environment.

### K562 culture

Human erythroleukemic cell line, K562 cells were cultured in RPMI-1640 medium (Gibco GranIsland, NY, USA) supplemented with 10% heat-inactivated fetal bovine serum (FBS; VWR), 1% L-glutamine (Gibco GranIsland, NY, USA) and 1% penicillin– streptomycin (Gibco). K562 cells were induced to differentiate using 3IU/ml recombinant human erythropoietin (EPO; PeproTech) following standard protocol.

### HUDEP-2 culture

Human umbilical-derived erythroid progenitor-2(HUDEP-2) were maintained in the culture as previously described[19]. HUDEP-2 cells were cultured in StemSpan™ Serum-Free Expansion Medium II (SFEM II; STEMCELL Technologies, Vancouver, BC, Canada) supplemented with stem cell factor (SCF; 50 ng/mL, PeproTech, Rocky Hill, NJ, USA), 3IU/ml erythropoietin (EPO; PeproTech), 1 μM dexamethasone (DEX; Sigma-Aldrich, St. Louis, MO, USA), 1 μg/ml doxycycline (DOX; Sigma-Aldrich) and 2% penicillin–streptomycin.

To induce differentiation, 2-5 million HUDEP-2 cells were grown in 1X Iscove’s modified Dulbecco’s medium, IMDM(Gibco) supplemented with 2% fetal bovine serum (FBS), 3% human blood type AB serum (Sigma-Aldrich),3IU/ml EPO, 10µg/ml Insulin (I9278, Sigma-Aldrich), 2IU/ml heparin(Sigma-Aldrich), 500µg/ml holo-transferrin (Sigma-Aldrich), 1% penicillin–streptomycin, 1% L-Glutamine and 1 μg/ml doxycycline. Media was replaced on day 4 and cells were maintained for another 2-3 days before analysis.

### Ex vivo expansion of primary human erythroblasts

Human peripheral blood samples were obtained from HbE/β-thalassemia patients at Nil Ratan Sircar Medical College and Hospital (NRSMCH), Kolkata. CD34+ HSPCs from peripheral blood samples were subsequently isolated using MACS CD34+ MicroBead kit (Miltenyi Biotec) according to manufacturer’s instructions. For ex vivo expansion, cells were cultured in proliferation media containing SFEM II supplemented with 100ng/ml SCF, 50ng/ml FLT3-L(PeproTech), 50ng/ml thrombopoietin (TPO; PeproTech), 20ng/ml IL-3(Miltenyi Biotec) and 1% penicillin–streptomycin. To induce proliferation and differentiation, human CD34+ HSPCs were cultured in two-phase culture system. In phase 1, cells were maintained in IMDM medium supplemented with 100ng/ml hSCF, 2% penicillin–streptomycin, 1ng/ml IL-3, 330μg/ml of holotransferrin, 10μg/ml of insulin, 5% human A/B plasma, 3IU/ml EPO, and 10 μg/ml of heparin for 9 days. In phase 2, IL-3 was withdrawn and cells were cultured for another 3-4 days.

### HEK293 cell culture

HEK293 cells were cultured in Dulbecco’s modified Eagle’sMedium, DMEM (Gibco,GranIsland,NY, USA) supplemented with 10% FBS and 1 % penicillin– streptomycin–glutamine. Cells were lifted for passaging by using 0.05% trypsin–EDTA (Gibco, GranIsland, NY, USA) for 5 min at 37^0^C.

### LncRNA expression from Affymetrix HG-U133 plus 2.0 Array

We collected lncRNA annotations from two resources; Reference sequence (RefSeq) database and Ensembl database, to reannotate the array probe sets of the Affymetrix HG-U133 Plus 2.0 Array, and then mapped the probe sets to the human genome for extracting LncRNA expression as described previously by Liu et al[20]. The details regarding datasets recruitment and preprocessing were described in the Supplemental information (Figure S1).Probes with no mismatch and mapped uniquely to the reference genome were considered for further processing. After removing the pseudogenes transcripts or protein-coding transcripts, we obtained, at least, 2248 unique LncRNAs that mapped uniquely to their exons. We further performed the Significance Analysis of Microarrays (SAM) using the R package to evaluate the differentially expressed lncRNAs between the study groups (high HbF vs. normal HbF).The differentially expressed lncRNAs were imported in Cluster view 3.0to carry out the Hierarchical Cluster Analysis (HCA)[21].

### Identification of gene modules and hub LncRNAs using weighted gene co-expression network analysis (WGCNA)

We performed weighted gene co-expression network analysis(WGCNA) packages in R, using standard procedures[22]. WGCNA is an approach to utilize gene expression data to construct co-expression-based weighted correlations, and can be used to measure the correlation between lncRNAs and mRNAs in the datasets. This will help us to identify hub lncRNAs which can be crucial in fetal hemoglobin regulation. LncRNA expression profiles for co-expression analysis were obtained from GSE13284 dataset. Briefly, we have calculated Pearson’s correlation matrix for all pair-wise lncRNAs and subsequently, weighted adjacency matrix using the soft-thresholding parameters calculated by pickSoftThreshold function. Then, adjacency matrix has been transformed into topological overlap matrix (TOM) to represent the connectivity of each lncRNA and genes in the module. Hierarchical analysis dendrogram was also performed to merge the genes with similar co-expression pattern into modules. The networks were visualized using the Cytoscape software (version 3.7.0)[23].

### Drug treatment

To determine the optimal concentration of hydroxyurea (Sigma-Aldrich) treatment for efficient gamma globin induction in K562 cells, we aimed to treat the cells with a range of hydroxyurea concentrations. Erythroid differentiation was performed in presence of Recombinant human EPO (3IU/ml) for 72h. Post differentiation, flow cytometry was carried out to check the expression of CD71 and CD235a (Glycophorin-A). Differentiated K562 cells (2×10^5^) were treated with different concentrations of the drug (50-300µM) for 24h. MTT assay (Sigma-Aldrich) was performed to check the viability and proliferation of the cells post drug treatment.

### Giemsa-Wright staining

For morphological analysis, cells were harvested and smeared on glass slide by cytospin centrifugation. After air drying for 5 min, slides were stained with May-Grunwald (S-039, Himedia Laboratories, Mumbai, India) solution for 5 min, rinsed in 40mM Tris buffer (pH 7.2) for 2min and counterstained with 1:20 diluted Giemsa (S011, Himedia Laboratories) solution for 15 min. Stained cells were analysed, and images were taken using Nikon TE2000 microscope equipped with a digital camera.

### RNA isolation and RT-qPCR

Total RNA was extracted using the miRneasy mini kit (QIAGEN) according to the manufacturer’s instructions. For mRNA, Reverse transcription was performed by High-Capacity cDNA Reverse Transcription Kit (Applied Biosystems™, Foster City, CA, USA). PowerUp™ SYBR™ Green Master Mix (Applied Biosystems™) was used to perform qPCR. For miRNA expression studies, assays were carried out using miRCURY LNA universal RT miRNA PCR method (QIAGEN). For mRNAs and lncRNAs, the data were normalized using the endogenous β-actin control. Normalization of miRNA expression was performed using U6 snRNA. All Real-time PCR reactions were performed using QuantStudio 5 real-tme PCR systems (Applied Biosystems™). The reactions were performed in at least three different samples in triplicate. Data analysis was carried out using the ΔΔCT method. The primer sequences for RT-qPCR are enlisted in the Supplemental information (Table S5).

### Nucleus-cytoplasm fractionation

Here, K562 cells (2x10^7^) were harvested and washed with ice-cold PBS for subcellular fractionation. Cells were resuspended in ice-cold cytoplasmic lysis buffer**[0.05%(v/v) NP-40,0.5mM DTT,1mM EDTA, HEPES (pH7.9), 10mM KCL and 1.5mM MgCl_2_]**and incubated for 5 min on ice. Cell lysates were then spun at 16,000 g for 10 min at 4 °C and supernatant was collected as cytoplasmic fraction. The pellet containing the nuclei was washed with nuclei wash buffer (0.1% (v/v) Triton X-100, 1 mM EDTA, 100 U Ribolock RNase inhibitor). The final nuclear pellet was resuspended in **nuclei lysis buffer[26%(v/v) Glycerol, 0.2mM EDTA, 0.5mM DTT,HEPES (pH7.9), 300mM NaCl and 1.5mM MgCl_2_]** and incubated for 5 min on ice. The supernant was collected as nuclear fraction. RNA was extracted from cellular fractionation using Trizol method. RT-qPCR was performed to measure the abundance UCA1-lncRNA in subcellular fractions. The expression of Malat1 and Beta-actin were regarded as loading control for nuclear RNA and cytoplasmic RNA, respectively.

### miRNA expression profiling

Total RNA was isolated using miRNeasy Mini Kit (QIAGEN), and was reverse-transcribed using miScript II RT Kit (QIAGEN) according to the manufacturer’s protocol. The miScript miRNA PCR Array (QIAGEN, GmbH, Hilden, Germany) in 96 well formats were used for real-time PCR. For expression profiling, 2× QuantiTect SYBR Green PCR Master mix, 10× miScript Universal Primer, RNase free water and template cDNA were mixed according to manufacturer’s protocol. Cycling conditions for real-time PCR were: initial activation step at 95^0^C for 15 min, followed by 40 cycles of 3-step cycling (denaturation at 94^0^C for 15 s, annealing at 55^0^C for 30 s and extension at 70^0^C for 30 s). To verify specificity and identity, dissociation curve analysis was performed. RT-qPCR data was normalized using RNU6-6P. For quality check, PCR array reproducibility and RT efficiency tests were performed using GeneGlobe.

### miRNA target prediction

miRNA sequences, annotations and distributions were obtained from miRBase database[24]. The target sites of the miRNA were identified by using multiMiR R package and database[25], a comprehensive collection of predicted and validated miRNA target prediction database which compiled data from 14 different databases.

For lncRNA transcripts, miRNA target sites were predicted by using DIANA-LncBase v3.0 (www.microrna.gr/LncBase)[26]. All interactions were visualized by using RNAhybrid[27].

### Cell transfection

Erythroid cells were transfected using Lipofectamine™ 2000 reagent (Thermo Fisher Scientific) according to the manufacturer’s instructions. The transfections included miR-148b mimic, anti-miR-148b, and a scramble control (each at a final concentration of 100 nM), as well as siRNA targeting UCA1 (final concentration 50 nM). The DsiRNA sequences targeting UCA1 were designed using the DsiRNA Design Tool (Integrated DNA Technologies, Coralville, IA, USA) and synthesized by Eurofins Genomics Services. Cells were seeded in 12-well plates and transfected with either miR-148b mimic, anti-miR-148b, scramble control (QIAGEN), or si-UCA1. After 48h, cells were harvested for immunoblotting and RT-qPCR analyses. All experiments were independently repeated at least three times for statistical validation. The sequence of siRNAs are enlisted in the Supplemental information (Table S6).

### CellTiter-Glo® Cell Viability Assay

To check the cell viability after transfection,cells were harvested from 12-well plate and seeded in 96-well opaque culture plate (Corning, NY, USA). CellTiter-Glo® reagent (Promega, Madison, WI, USA) was added to each well and the luminescence signal was read after 15 minutes with GloMax® Navigator Microplate Luminometer (Promega).

### Stable transfection of UCA1-lncRNA

The full-length cDNA(1.4kb) of UCA1 was amplified using primers (forward: 5’-CGGGATCCTGACATTCTTCTGGACAAT-3’,reverse: 5’-GGAATTCGGCATATTAGCTTTAATGTAGG-3’). The PCR product was digested with BamHI and EcoRI restriction enzymes and subcloned into the pcDNA3.1-mcherry (a gift from David Bartel, Addgene plasmid # 128744)[28] and confirmed by sequencing. Thereafter, pcDNA3.1-mcherry/UCA1 construct was transfected into K562 cells using Lipofectamine™ 2000((Invitrogen, CA, USA) and screened with 1000µg/ml G418 disulfide (Sigma-Aldrich) for over 7 days. Transfection with pcDNA3.1-mcherry empty vector (EV) acted as control. The positive clones were confirmed by RT-qPCR for UCA1-lncRNA expression. The primers for UCA1-lncRNA were as follows: forward, 5’-GCCGAGAGCCGATCAGACAAA-3’; reverse, 5’-GCTGGGATGGCCATTTGGAAG-3’. Annealing temperature was at 59^0^ C.

### Immunofluorescence

For immunofluorescence study, 0.1 million K562 cells were collected in 1X PBS and spun on slides using CS II centrifuge (SLEE, Germany). Cells were washed with ice-cold PBS, and fixed using 0.05% glutaraldehyde (Sigma-Aldrich) for 10 min. After washing with 1X PBS twice, cells were permeabilized with 0.1% Triton-X for 5 min at room temperature. Thereafter, cells were incubated with specific primary antibody,Hemoglobin γ (D4K7X) Rabbit mAb (Cell Signaling Technology, Waltham, MA, USA; 1:500 dilution) overnight at 4^0^C followed by Alexa Fluor® 594-conjugated secondary antibody (1:2000,ab150080; Abcam, Cambridge, MA,USA) in the dark for 1 h at room temperature. The nuclear marker DAPI was added for 3-5 min. The cells were observed using fluorescent microscopy.

### Dual-luciferase reporter assay

To construct the luciferase reporter vectors, the human BCL11A, ZBTB7A gene 3’-UTR fragments and the sequence of UCA1-lncRNA containing the potential binding sites for miR-148b were inserted into SacⅠ/XbaⅠ site in pmirGLO Dual - Luciferase miRNA Target Expression Vector (Promega) downstream of the firefly luciferase gene. The target-site mutant vectors for each target were generated by replacing the target sequence with scramble sequence. The WT and mutant reporter vectors were transfected into HEK293T cells, along with miRCURY LNA miRNA Mimics, miR-148b mimics (QIAGEN)using a Lipofectamine 2000 Transfection Reagent. The expression Renilla luciferase gene was used as an internal reference for transfection efficiency. Cells were lysed 48h after transfection, and luciferase activity was analysed using the dual-luciferase reporter assay system (Promega) according to the manufacturer’s instructions. All the experiments were independently repeated at least three times for the statistical validation.

### Flow cytometry

To evaluate erythroid differentiation, cells were washed twice with phosphate-buffered saline/0.5% BSA and fixed using 0.05 %(v/v)glutaraldehyde. Cells were incubated with PE-conjugated anti-CD71 and FITC-conjugated anti-CD235a antibodies (Miltenyi Biotec, Bergisch Gladbach, Germany) on ice for 30 min at dark. Thereafter, flow cytometry was performed.

For HbF staining, 1-2 million cells were washed with PBS/0.5% BSA and fixed using 0.05 %(v/v) glutaraldehyde for 10 min. After repeated washes with PBS/0.5% BSA, cells were permeabilized with 0.1% Triton for 5 min at room temperature. Cells were stained with APC conjugated HbF antibody (Miltenyi Biotec) on ice for 20-30 min. All data were acquired in a BD FACSCanto machine and analysed using FlowJo™ software (FlowJo LLC, Ashland, OR, USA).

### Apoptosis and cell cycle analysis

Transfected cells were harvested after transfection and double stained with fluorescein isothiocyanate (FITC)-annexin V and propidium iodide (PI) using FITC Annexin V/Dead Cell Apoptosis Kit (Invitrogen) according to the manufacturer’s instructions. Cells were discriminated into viable cells, dead cells, early apoptotic cells and late apoptotic cells. The percentage of different cells were counted and compared. For cell cycle analysis, cells were stained with propidium iodide (PI) and analysed by flow cytometry. The percentage of the cells in G0-G1, S, and G2M phases were calculated and compared. All analyses were performed using FlowJo™ Software.

### Immunoblotting

Cells were collected and lysed with RIPA buffer (50 mM Tris-HCl, pH 7.4, 150 mM NaCl, 1% Triton X-100, 0.5% sodium deoxycholate, 0.1% SDS, 1 mM EDTA) following standard protocol. Equivalent amounts of protein (10-20µg) were run on the 12% SDS–polyacrylamide gel, and transferred to nitrocellulose membrane. Immunoblotting of the membranes was performed using the following primary antibodies: 1:1000; anti-gamma globin (Cell Signaling Technology), 1:1000; anti-BCL11A (A5445;Abclonal, Wuhan, China), 1:1000; anti-β-Actin (ab8227, Abcam). Signals were evaluated after incubation with appropriate secondary antibody coupled with horseradish peroxidase (HRP). Scanned images were quantified using ImageJ software.

### RP-HPLC

The globin chain analysis of the differentiated cells were performed as previously described[29]. The differentiated cells (0.5-1 million) were washed twice with PBS and lysed using 100µl HPLC-grade water (SRL chemicals, Mumbai, India). Cell debris was cleared by centrifugation at maximum speed on a desktop centrifuge. The supernatant (90µl) was taken and mixed with 10µl Tris(2-carboxyethyl) phosphine hydrochloride (TCEP.HCl) to break the disulfide bonds. After 5 min incubation at room temperature, 85µl of 0.1% trifluoroacetic acid(TFA)/32% acetonitrile were added and vortexed briefly. A 50µl aliquot of the solution was analysed at 0.7 ml/min flow rate for 50 min using Agilent 1260 Infinity II LC System. Commercially available standard hemoglobin containing HbA, HbD, HbF etc. were used as reference isotypes. The globin chains were detected at 215nm.

### RNA–RNA pull-down assay

The UCA1-overexpressing K562 cells were transfected with 100 pmol of biotinylated miR-148b mimics or non-specific control using Lipofectamine 2000(Thermo Fisher Scientific, Waltham, MA, USA) according to the manufacturer’s instructions. Twenty-four hours post-transfection, cells were cross-linked with 1% paraformaldehyde in PBS for 10 min at room temperature (RT) under gentle agitation. Cross-linking was quenched by the addition of glycine to a final concentration of 0.125 M, followed by two washes with ice-cold PBS. Cells were collected in PBS, pelleted at 510 g for 5 min at 4 °C, and lysed in lysis buffer containing **50 mM Tris-HCl (pH 7.0), 10 mM EDTA, 1% SDS, RNase inhibitor (200 U/mL), and protease inhibitors (5 μL/mL)**. Lysates were incubated with streptavidin magnetic beads overnight at RT under moderate agitation to isolate RNA complexes associated with biotinylated miRNAs. Beads were washed five times with wash buffer containing 0.5% SDS in **2× SSC** at RT, each with 5 min agitation. Bound complexes were digested with proteinase K (1 mg/mL) in buffer (**10 mM Tris-HCl, 100 mM NaCl, 1 mM EDTA, 0.5% SDS**) for 45 min at 50 °C, followed by 10 min at 95 °C to reverse cross-links. RNA was purified using a commercial RNA purification kit including on-column DNase digestion, and RT-qPCR was performed using UCA1-specific primers.

### Statistical analysis

All results were represented as Mean ± SD. Individual data points representing biological replicates were included in the graph. All experiments were repeated at least three times. The comparison between two groups was performed by unpaired two-tailed t-test. Differences among more than two groups were evaluated using one-way ANOVA, followed by Bonferroni post-hoc analyses as appropriate. All analyses were carried out using GraphPad software 8.0.p <0.05 was considered statistically significant.

## ETHICAL APPROVAL

The ethical approval for the study was obtained from IIT Kharagpur and NRS Medical College, Kolkata prior to sample collection. (Approval Nos. IIT/SRIC/DR/2019 dated 06.11.2019, IIT/SRIC/DEAN/2022 dated 12.08.2022; No/NMC/3483 dated 12.07.2019; IIT/SRIC/SAO/2017 dated 11.12.2017; No/NMC/4154 dated 03.08.2016No/NMC/3483, No/NMC/4154 and IIT/SRIC/SAO/2017).

## DATA SHARING STATEMENT

The data are not publicly available due to privacy or ethical restrictions. For original data, please.

## AUTHOR CONTRIBUTIONS STATEMENT

M.R. and N.C. conceptualized the study. M.R. and C.B. performed the in silico analyses. M.R., S.B., and M.M. coordinated the blood sample collection, which was carried out by trained clinical personnel. M.R., S.B., and M.M. were involved in data collection, curation, and analysis. M.R. and S.B. prepared the figure panels and wrote the manuscript. N.C., P.C.S., and T.K.D. reviewed and edited the manuscript. All authors have read and approved the final version of the manuscript.

## ACKNOWLEDGEMENTS

The authors would like to acknowledge the infrastructural supports and services provided by the School of Medical Science and Technology, Indian Institute of Technology Kharagpur and NRS Medical College and Hospital, Kolkata. The authors would like to thank Dr. Ryo Kurita and Dr. Yukio Nakamura from the Cell Engineering Division, RIKEN BioResource Research Center, for generously providing the HUDEP-2 cells. The authors also acknowledge Dr. Sivaprakash Ramalingam from CSIR-Institute of Genomics and Integrative Biology (CSIR-IGIB) for providing the K562 cells. The authors would like to extend special thanks to the members of Dr. Gayatri Mukherjee’s lab at the School of Medical Science and Technology (SMST), IIT Kharagpur, for their valuable support and assistance with flow cytometry. The authors acknowledge the use of OpenAI’s ChatGPT, a large language model, for assistance in language editing and improving the clarity of the manuscript. All content was reviewed and verified by the authors. Mr. Motiur Rahaman would like to acknowledge the Ministry of Education for financial support. This work is supported by the Department of Biotechnology (DBT), Ministry of Science and Technology, Government of India. The project is titled as “MicroRNA based reprogramming of fetal hemoglobin in β-thalassemia” [Sanction no: BT/PR32054/MED/97/454/2019].

## COMPETING INTERESTS

The authors have no conflict of interest to declare.

## Notes

### Competing Interest Statement

The authors have declared no competing interest.

