## Supplementary Files for "UCA1 lncRNA represses γ-globin expression by sequestering miR-148b, a key post-transcriptional regulator of BCL11A"

### Supplementary data

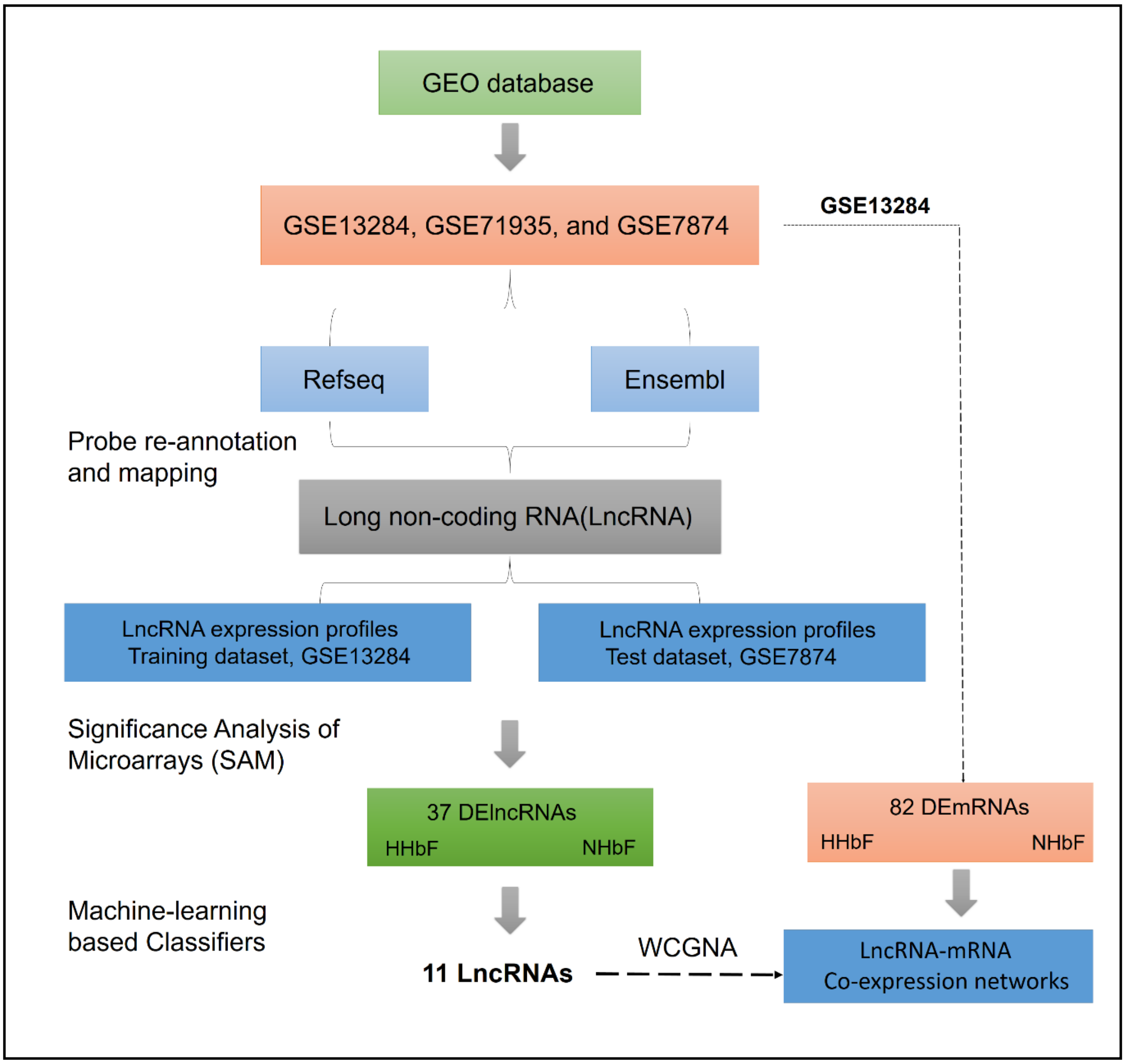

**Figure S1.** Schematic representation of the in silico pipeline used to develop lncRNA-based classifiers for identifying high fetal hemoglobin (HbF) conditions. The workflow includes data acquisition, preprocessing, differential expression analysis, feature selection, model training using machine learning algorithms, and classifier evaluation.

**Table S1. Phenotypic and genotypic features of six unrelated HbEβ-thalassemia patients with high (HHbF) or normal (NHbF) fetal hemoglobin levels.**

| **Characteristics** | **HHBF1** | **HHBF2** | **HHBF3** | **NHBF1** | **NHBF2** | **NHBF3** |
|---|---|---|---|---|---|---|
| Age(years)/sex | 6/F | 15/M | 9/F | 15/F | 6/M | 16/M |
| Age of onset(years) | 2.5 | 2 | 3 | 3 | 2 | 1.5 |
| No. of transfusion/year | 0 | 0 | 0 | 12 | 12 | 8 |
| Hb(g/dL) | 7.8 | 7.6 | 8.2 | 6.5 | 7.2 | 7.4 |
| HbF(%) | 52.5 | 47.7 | 45.3 | 5.6 | 3.1 | 6.9 |
| HbA0 | 0.5 | 0 | 0 | 26.9 | 32.1 | 12 |
| HbA2+E (%) | 45.7 | 35.5 | 53.2 | 57.2 | 52.5 | 75.2 |
| MCV(fL) | 67 | 65.4 | 66 | 70.2 | 68.9 | 72 |
| RDW | 29.8 | 28.6 | 28.9 | 29.8 | 30.9 | 35.6 |
| MCH(g/dL) | 20.5 | 20 | 19.8 | 20.8 | 21.4 | 22.5 |
| **HBB genotype*** | IVS-1-5(G>C)/cd26 | IVS-1-5(G>C)/cd26 | IVS-1-5(G>C)/cd26 | IVS-1-5(G>C)/cd26 | IVS-1-5(G>C)/cd26 | IVS-1-5(G>C)/cd26 |

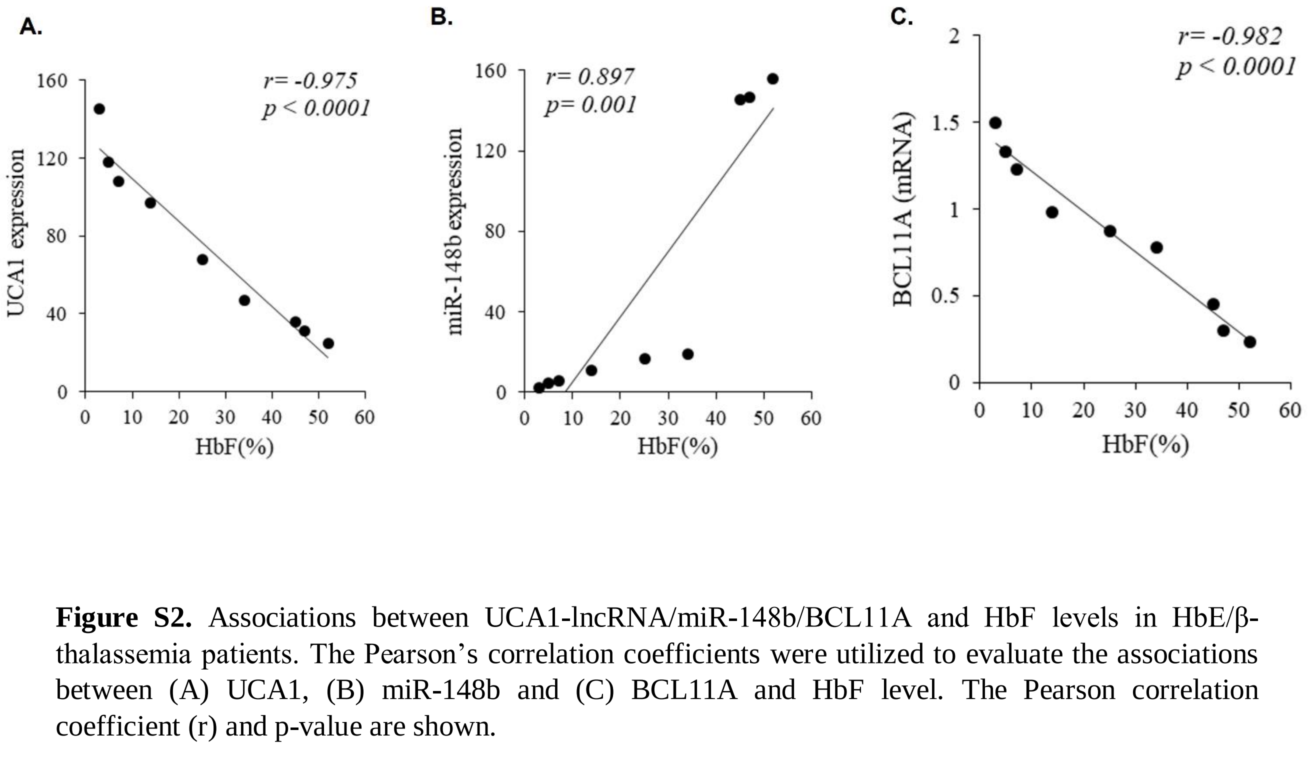

**Figure S2.** Associations between UCA1-lncRNA/miR-148b/BCL11A and HbF levels in HbE/β-thalassemia patients. The Pearson's correlation coefficients were utilized to evaluate the associations between (A) UCA1, (B) miR-148b and (C) BCL11A and HbF level. The Pearson correlation coefficient (r) and p-value are shown.

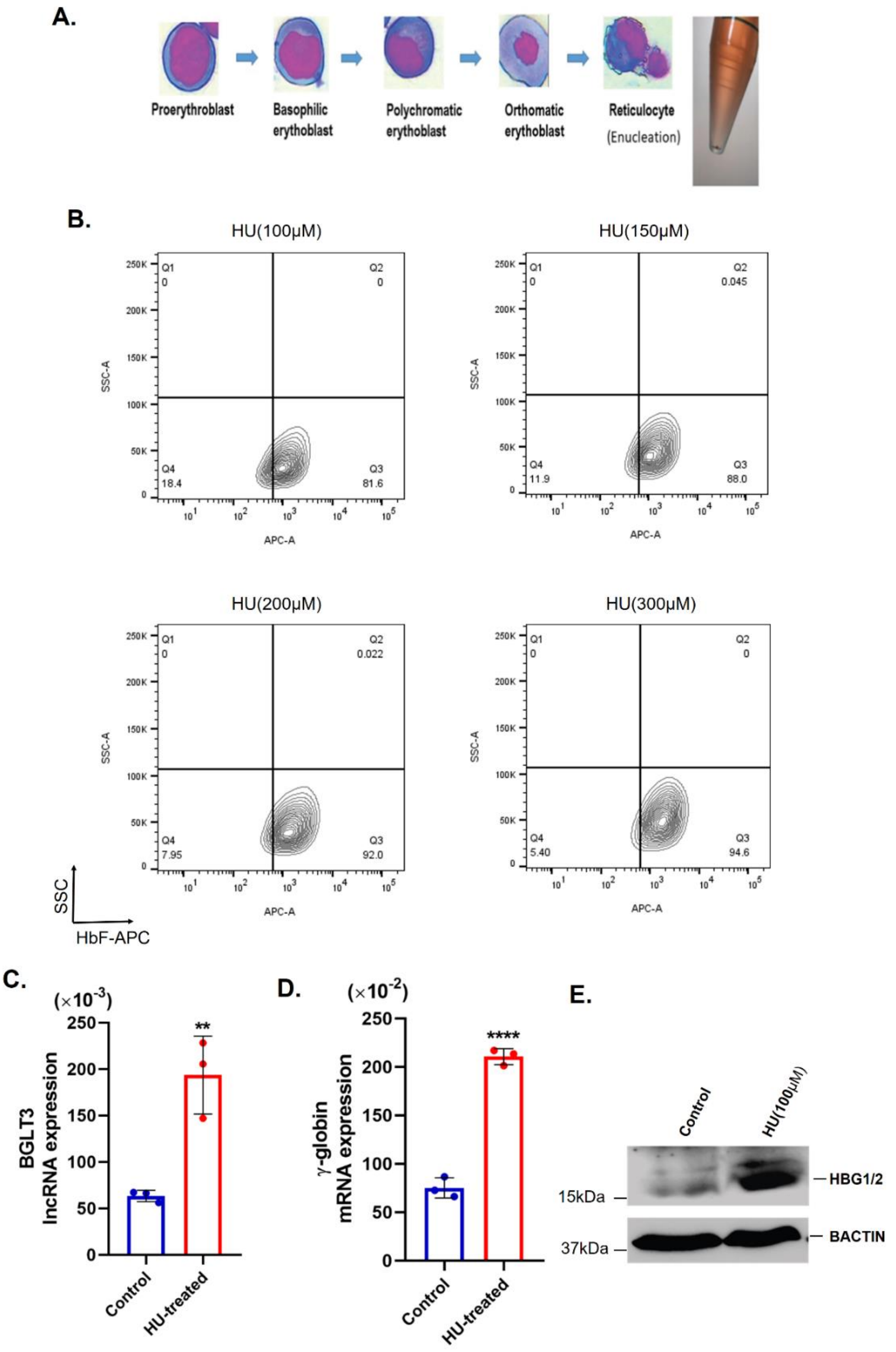

**Figure S3.** (A) Kinetics of erythroid differentiation of K562 cells determined by Giemsa staining after culturing in EPO (3IU/ml) containing media. (B) Representative F-cell staining flow cytometry plots of K562 cells treated with different concentrations (100-300µM) of Hydroxyurea (HU). C. Gene expression of (C) BGLT3 and (D) Gamma globin in control and HU-treated K562 cells determined by qRT-PCR. (E) Western Blot analysis with indicated antibodies of cell lysates from control and HU-treated K562 cells. Beta-actin served as a loading control. Data are shown as mean ± SD of three independent experiments. Two-tailed Student's t-test, *p<0.05, ***p<0.01, ****p< 0.001.

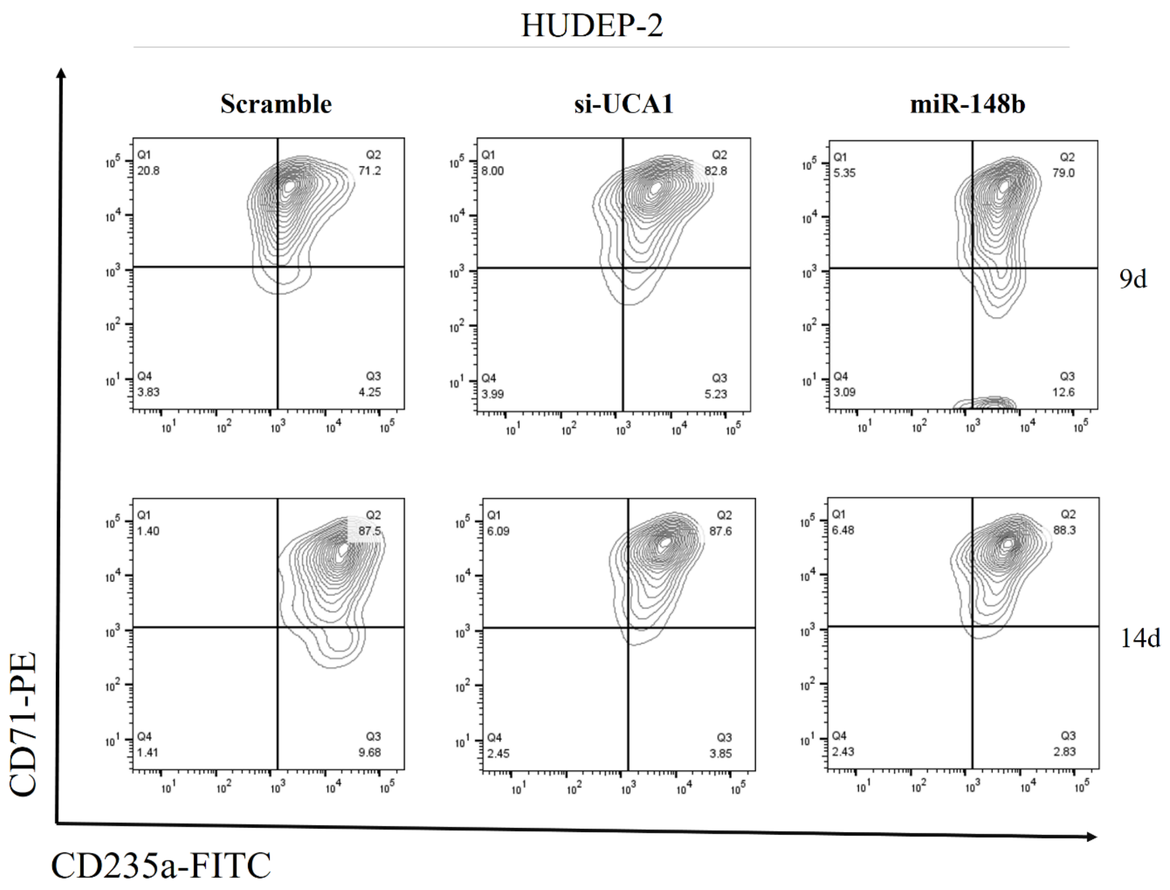

**Figure S4.** Kinetics of erythroid maturation of scramble, si-UCA1 and miR-148b transfected HUDEP-2 cells determined by flow cytometry for CD71 and CD235a at the indicated time points after culture in erythroid differentiation medium.

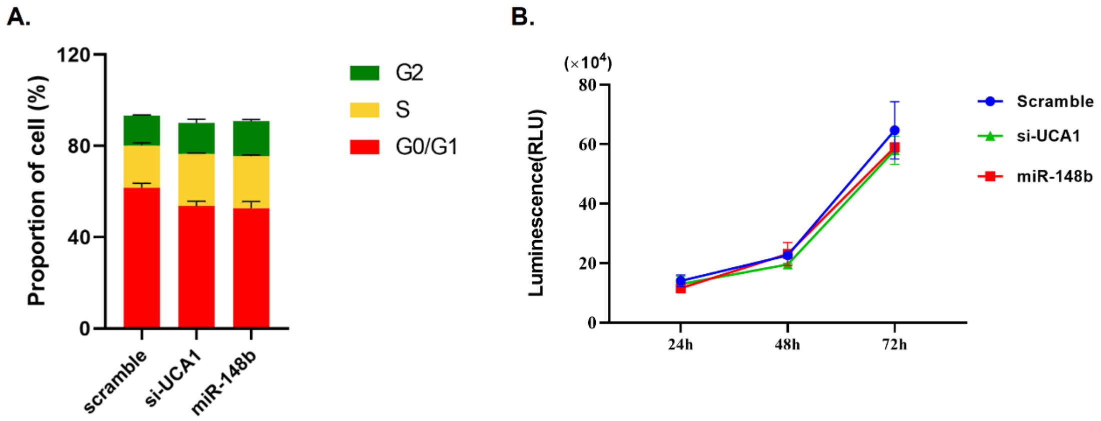

**Figure S5.** A. Cell cycle assay of si-UCA1, miR-148b and scrambled control transfected HUDEP-2. B. Proliferation assay of si-UCA1, miR-148b and scrambled control HUDEP-2 cells. CellTiterGLO® Luminescent Cell Viability assay method.

**Table S2. List of differentially expressed lncRNAsidentified from normalized data using the Significance Analysis of Microarrays (SAM).**

| **Significant Positive** | | | |
|---|---|---|---|
| **Row** | **Gene ID** | **Gene Name** | **Score(d)** |
| 95 | 1554715_at | DRAIC | 2.0437013764306 |
| 1162 | 222090_at | NDUFB2-AS1 | 1.68538370889437 |
| 1789 | 240115_at | PPM1F-AS1 | 1.67859542171533 |
| 28 | 1553086_at | C11orf40 | 1.63136538695405 |
| 20 | 1552954_at | LINC02899 | 1.62937369394437 |
| 716 | 1562597_at | LINC02150 | 1.58376206957499 |
| 1908 | 242987_x_at | LAMA5-AS1 | 1.55079093882051 |
| 1103 | 216596_at | LINC02249 | 1.54514911414644 |
| 1101 | 216473_x_at | DBET | 1.5138023689809 |
| 81 | 1553935_at | BFSP2-AS1 | 1.48864525059644 |
| 708 | 1562478_at | LINC00659 | 1.47046939273058 |
| 1537 | 234293_x_at | RAB4A-AS1 | 1.46118196197715 |
| 1349 | 230432_at | LINC02532 | 1.4518654546811 |
| 1875 | 241743_at | ZBTB47-AS1 | 1.41845679914795 |
| 1508 | 233238_s_at | LINC01933 | 1.41715312017631 |
| 514 | 1560550_at | LINC01644 | 1.4165684639935 |
| 1369 | 230743_at | HOXB-AS3 | 1.39012580215454 |
| 134 | 1555994_at | DIAPH3-AS1 | 1.38923460999856 |
| 1492 | 232897_at | LERFS | 1.38535652775818 |
| 1472 | 232571_at | CAPN10-DT | 1.38186863924982 |
| **Significant Negative** | | | |
| **Row** | **Gene ID** | **Gene Name** | **Score(d)** |
| 1381 | 230944_at | C6orf223 | -2.38780262879588 |
| 1246 | 227925_at | GSEC | -1.79483433782607 |
| 1245 | 227919_at | UCA1 | -1.71932097321119 |
| 160 | 1556406_at | LINC00879 | -1.71603145040091 |
| 1201 | 225457_s_at | PP7080 | -1.70201243115296 |
| 1741 | 239113_at | CT66 | -1.68085518235983 |
| 1368 | 230710_at | MIR210HG | -1.66092713399978 |
| 1290 | 229090_at | ZEB1-AS1 | -1.65242417165371 |
| 1446 | 231954_at | MIR4453HG | -1.64110553679087 |
| 1699 | 238021_s_at | CRNDE | -1.60334312036694 |
| 1280 | 228839_s_at | LINC00863 | -1.59751875990652 |
| 989 | 1569807_at | LINC01741 | -1.54994192087381 |
| 726 | 1562655_at | SCUBE1-AS1 | -1.50779860571192 |
| 649 | 1561565_at | LINC02530 | -1.46763779701633 |
| 1090 | 215590_x_at | ACVR2B-AS1 | -1.44870642262329 |
| 185 | 1556677_at | ZFY-AS1 | -1.42882118868754 |
| 1450 | 232239_at | LINC00865 | -1.42237920007859 |

**Table S3. The five most significantly up- and down-regulated miRNAs in patient CD34+ cells**

| Mature ID | Fold Regulation | p-value | miScriptCatalog |
|---|---|---|---|
| **Up-Regulation** | | | |
| hsa-miR-218-5p | 2.1594 | 0.015544 | MIMAT0000275 |
| hsa-miR-148b-3p | 1.8204 | 0.027273 | MIMAT0000759 |
| hsa-miR-146a-5p | 1.2652 | 0.01769 | MIMAT0000449 |
| hsa-miR-26a-5p | 1.0144 | 0.000831 | MIMAT0000082 |
| hsa-miR-125b-5p | 1.0028 | 0.013907 | MIMAT0000423 |
| **Down-Regulation** | | | |
| hsa-miR-320a | -2.03 | 0.042007 | MIMAT0000510 |
| hsa-miR-302c-3p | -1.8311 | 0.022462 | MIMAT0000717 |
| hsa-let-7b-5p | -1.0688 | 0.023121 | MS00003122 |
| hsa-miR-15b-5p | -1.0229 | 0.004408 | MIMAT0000417 |
| hsa-miR-92a-3p | -1.0004 | 0.012087 | MIMAT0000092 |

**Table S4. The five most significantly up- and down-regulated miRNAs in K562 cells**

| Mature ID | Fold Regulation | p-value | miScriptCatalog |
|---|---|---|---|
| **Up-Regulation** | | | |
| hsa-miR-98-5p | 4.6 | 0.00087 | MIMAT0000096 |
| hsa-miR-375 | 3.96 | 0.009343594 | MS00004088 |
| hsa-miR-148b-3p | 3.39 | 0.0303 | MIMAT0000759 |
| hsa-miR-185-5p | 3.34 | 0.000488049 | MS00003647 |
| hsa-miR-124-3p | 3.3 | 0.002721251 | MS00006622 |
| **Down-Regulation** | | | |
| hsa-miR-92a-3p | -3 | 0.0043 | MIMAT0000092 |
| hsa-let-7b-5p | -3.03 | 0.003802762 | MS00003122 |
| hsa-miR-320a | -3.03 | 0.0087 | MIMAT0000510 |
| hsa-miR-7-5p | -3.11 | 0.000680456 | MS00006503 |
| hsa-miR-142-3p | -3.66 | 0.010521821 | MS00006664 |

A.

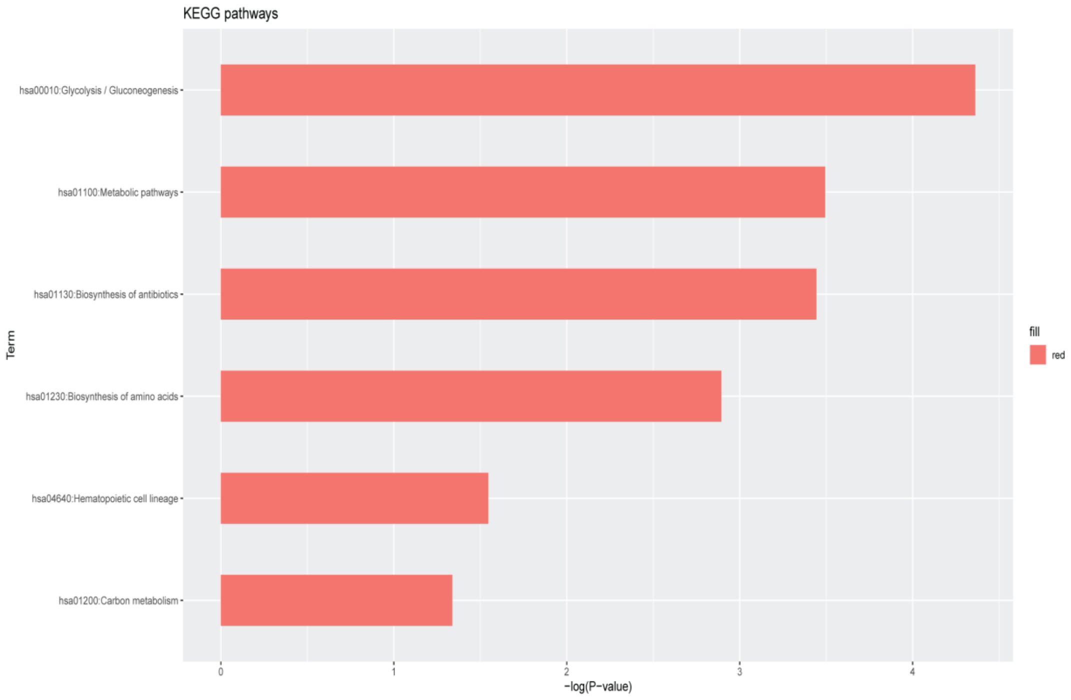

B.

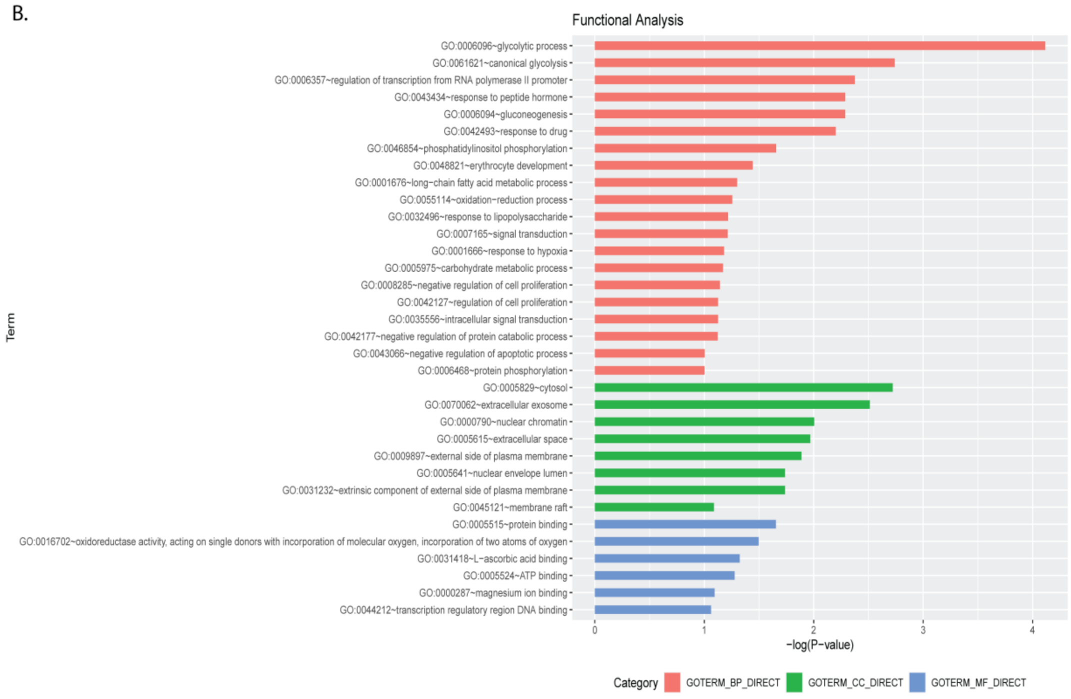

**Figure S6.**GO term and functional enrichment analysis. (A) KEGG pathway analysis were performed in differentially expressed mRNAs correlated with 11 lncRNAs. The 6 significant ($p<0.05$) signaling pathways are shown. All values are negative log transformed.(B) Gene Ontology (GO) annotation of differentially expressed mRNAs correlated with 11 lncRNAs, with a list top enriched genes ($p<0.05$), covering the domain of biological processes, cellular components and molecular functions. Enrichment values were –log10 (p-value) transformed**.**

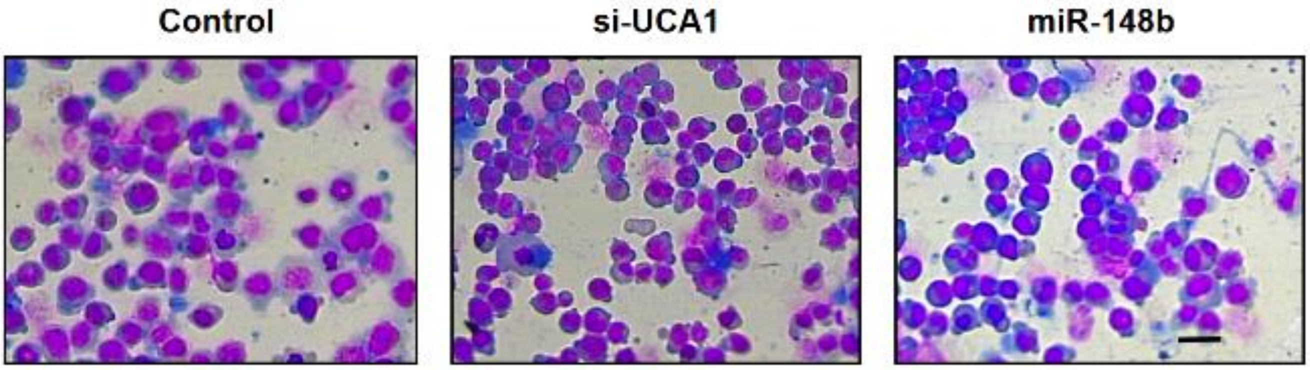

**Figure S7**. Representative images of Wright-Giemsa-stained cytospin of CD34+ HSPCs: Scramble control (Left), si-UCA1 (middle) and miR-148b (right). Scale bars, 50μM.

**Table S5. List of Primers for PCR and qRT-PCR**

| Gene name | NCBI Gene Id | Primer sequence (5' > 3') |
|---|---|---|
| **Endogenous Control genes(qRT-PCR)** | | |
| Beta-actin(Cytoplasmic Fraction) | X00351.1 | F-ACTGGAACGGTGAAGGTGACA |
| | | R-AGTCCTCGGCCACATTGTGAA |
| Malat1(Nuclear Fraction) | NR_002819.5 | F-AGCAAACTGTGTTGGCGTGG |
| | | R-CGGTGCCTTTAGTGAGGGGT |
| **Globin genes(qRT-PCR)** | | |
| HBG | NM_000184.3 | F-AAGCTCCTAGTCCAGACGCC |
| | | R-AGACAACCAGGAGCCTTCCC |
| HBB | NM_000518.4 | F-TGGATGAAGTTGGTGGTGAG |
| | | R-CCTTAGGGTTGCCCATAACA |
| HBA1 | NM_000558.5 | F-CCCGGTCAACTTCAAGCTCCTA |
| | | R-AAGAAGCATGGCCACCGAGG |
| HBA2 | NM_000517.6 | F-CCGGTCAACTTCAAGCTCCTA |
| | | R-AGGAGGAACGGCTACCGAG |
| **miRNA target genes(qRT-PCR)** | | |
| BCL11A | NM_001405730.1 | F-GACGCAGCGACACTTGTTCT |
| | | R-GCTCTCGAGCTTCCATCCGA |
| ZBTB7A | NM_015898.4 | F-ATCCGAGCCAAGGCCTTCCA |
| | | R-ACCTTCAGCTTGTCCTGCCTGG |
| **LncRNA Primers(qRT-PCR)** | | |
| ZEB1-AS1 | NR_024284.1 | F-CTACGGCCGGAACCTTGTTG |
| | | R-AAACCAGGCGTCCCTTTCCAA |
| UCA1 | NR_015379.3 | F-GCCGAGAGCCGATCAGACAAA |
| | | R-GCTGGGATGGCCATTTGGAAG |
| GSEC | NR_033839.1 | F-GCCTGATGGGGATACCTTCC |
| | | R-CAAGGCCAGGGTTTAGGTGA |
| MIR4453HG | NR_033797.2 | F-TCCCCCTAGCCATGAAAGGA |
| | | R-CTTGCTCGGCATTTCGTCTC |

| | | |
|---|---|---|
| BGL3 | NR_121648.1 | F-CAGGGGTAACACACAAACCAGC |
| | | R-CACACTTCCACCGGCAGAGA |
| **Erythroid differentiation markers(qRT-PCR)** | | |
| GATA1 | NM_002049.4 | F-GCCACTACCTATGCAACGCC |
| | | R-CCCGTTTACTGACAATCAGGC |
| BAND3 | X12609.1 | F-AACGTAGCTGGTCGCAGAG |
| | | R-TGTCTACGGTGATCTGAGCC |
| ALAS2 | NM_000032.5 | F-AGGAAGCCATTTTCCGGTCC |
| | | R-ACTGAAGACATAGTTTCCAGGC |
| **Overexpression UCA1(PCR)** | | |
| OUCA1-F(BamH1) | NR_015379.3 | F-CGGGATCCTGACATTCTTCTGGACAAT |
| OUCA1-R(EcoR1) | | R-GGAATTCGGCATATTAGCTTTAATGTAGG |
| **Beta-globin sequencing primers(PCR)** | | |
| BetaSeq-F | | F-GCATATTCTGGAGACGCAGGAAG |
| BetaSeq-R | | R-CATCAAGGGTCCCATAGACTCACC |

**Table S6. List of oligonucleotide sequences used for cloning and knock-down experiments**

| Gene Name | Oligonucleotide sequence(5'>3') |
|---|---|
| BCL11A-F | CGCGGCCGCAGCACTGGGTGAGGTAATAAACCTTAGGAACTAT |
| BCL11A-R | CTAGATAGTTCCTAAGGTTTATTACCTCACCCAGTGCTGCGGCCGCGAGCT |
| MutBCL11A-F | CGCGGCCGC AGATGGGTGAGGTAATAAACCTTAGGAACTAT |
| MutBCL11A-R | CTAGATAGTTCCTAAGGTTTATTACCTCACCCATCTGCGGCCGCGAGCT |
| ZBTB7A-F | CGCGGCCGCGGCAGGAGCGGTCACCTCACAGGTGGTGACACTGAGT |
| ZBTB7A-R | CTAGACTCAGTGTCACCACCTGTGAGGTGACCGCTCCTGCCGCGGCCGCGAGCT |
| MutZBTB7A-F | CGCGGCCGCGGCAGGAGCGGTCACCTCACAGGTGGTGACAGAGT |
| MutZBTB7A-R | CTAGACTCTGTCACCACCTGTGAGGTGACCGCTCCTGCCGCGGCCGCGAGCT |
| UCA1-F | CCCTCTCCTATCTCCCTTCACTGACTCTCTTTTCGGACTCAT |
| UCA1-R | CTAGATGAGTCCGAAAAGAGAGTCAGTGAAGGGAGATAGGAGAGGGAGCT |
| MutUCA1-F | CCCTCTCCTATCTCCCTTCTTACTCTCTTTTCGGACTCAT |
| MutUCA1-R | CTAGATGAGTCCGAAAAGAGAGTAAGAAGGGAGATAGGAGAGGGAGCT |
| si-UCA1 | CUGCAAUCAGAACUAUUGAACUUCU |
| | AGAAGUUCAAUAGUUCUGAUUGCAGAU |
